# Interplay of genotypic and thermal sex determination shapes climatic distribution in herpetofauna

**DOI:** 10.1101/2024.04.21.589911

**Authors:** Edina Nemesházi, Veronika Bókony

## Abstract

Sex is a fundamental trait of all sexually reproducing organisms, and sex-determination systems show a great diversity across the tree of life. A growing body of evidence shows that genotypic and temperature-dependent sex determination (GSD and TSD, respectively) can coexist, which theoretically can have wide-ranging consequences for demography and population persistence, especially under climate change. Temperature-induced sex reversal, resulting from combined effects of sex chromosomes and environmental temperatures on sexual development, can explain the frequent transitions between GSD and TSD, and even between different GSD systems, that happened multiple times in ectothermic vertebrates. However, general lack of empirical data on the prevalence of sex reversal has long constrained the assessment of its evolutionary-ecological significance. Here we analysed an exhaustive compilation of available data to demonstrate that the climatic distribution of extant species is explained by the combination of their sex-chromosome system (GSD) and temperature reaction norm (TSD) across the phylogeny of amphibians and some reptiles. This pattern is in accordance with predictions of the ‘asymmetrical sex reversal’ theory, underscoring the importance of temperature-induced sex reversal in phylogeography, evolution, and species conservation under the threat of climate change, and highlighting the need for more empirical research on sex reversal in nature.

## Introduction

The great diversity of sex-determination systems across animal phylogenies has long been puzzling researchers. For example, while mammals and birds possess genotypic sex determination (GSD), where individuals develop gonads corresponding to their sex chromosomes, the picture is much more complex in ectothermic vertebrates, where besides GSD, temperature-dependent sex determination (TSD) is also widespread ^1,2^. The type of sex determination has crucial implications for a diverse array of biological phenomena, including genome evolution, adaptive radiation, and the evolution of life-history and demographic traits and reproductive systems ^3–11^. Climatic change is inferred to be a major driver of the evolution of sex-determination systems ^5,12^, so understanding the mechanisms and consequences of these evolutionary processes is critical under the light of contemporary rapid climatic changes.

Theoretical models indicate that changing environmental temperatures could facilitate transitions between GSD and TSD ^13–16^, as well as between male-heterogametic (XX/XY) and female-heterogametic (ZW/ZZ) GSD systems ^14^ _via_ the mechanism of thermal sex reversal. This means that environmental temperatures can override the sexual development otherwise determined by sex chromosomes (and/or other genomic elements) in GSD, leading to a mismatch between genotypic sex and phenotypic sex. Sex-reversed individuals have been reported from a few free-living populations of ectothermic vertebrates ^17–22^, and circumstantial data suggest that sex reversal may occur in many more species than currently known ^1,12,23^. Because sex reversal may lead to sex-ratio bias which may then influence the fitness of sex-reversed and sex-concordant individuals differently (sometimes in favour of the sex-reversed; ^24,25^), temperature-induced sex reversal may have both evolutionary significance and implications that could help identify vulnerable hotspots for biodiversity conservation under the concurrent climate change. However, due to the scarcity of empirical data on sex reversal, it remains a challenge to evaluate its role in the evolution of ecological and other species-level traits across the tree of life.

As a result of sex-chromosome evolution, the type of GSD system might influence the direction in which sex reversal could occur in a species. According to Muller’s ratchet model, deleterious mutations tend to accumulate on the hemizygous sex chromosome which always occurs in heterogametic form (i.e. the Y chromosome in XX/XY and the W chromosome in ZW/ZZ systems; ^26^). If heterogametic individuals undergo sex reversal, 25% of their offspring will possess this chromosome in a homogametic form (i.e. YY or WW genotypes resulting from mating of XY males and XY females or ZW males and ZW females, respectively). Production of this new, potentially infertile or lethal genotype is expected to reduce fitness in sex-reversed heterogametic individuals. By contrast, sex reversal may be neutral or even advantageous in the homogametic sex ^25,27,28^. Furthermore, sex-antagonistic genes ^29^ accumulating on the Y or W chromosome may also cause disadvantage in sex-reversed XY or ZW individuals. Therefore, natural selection should favour resistance to sex reversal in the heterogametic sex, leading to biased sex-reversal patterns where sex-reversing environmental conditions affect the homogametic sex more prominently^16,30^. With other words, female-to-male sex reversal should be more easily induced in genetic females (XX) in XX/XY systems, while in ZW/ZZ systems, genetic males (ZZ) should be more inclined to undergo sex reversal and become phenotypic females, leading to ‘asymmetrical sex reversal’ ^30^; a theoretical explanation of the empirical pattern described as “Witschi’s rule” ^31^.

The theory of asymmetrical sex reversal provides testable predictions for the interplay between genotypic and thermal sex determination under different climatic conditions (Fig. 1). Accordingly, when directional climate change results in a thermal environment which would induce sex reversal in heterogametic individuals, sex-reversal resistance is expected to evolve in response to fitness reduction due to the production of YY or WW offspring, and this resistance should then allow the prevailing sex-chromosome system to persist longer (Fig. 1). Therefore, the climatic distribution of extant species should be constrained by their sex-chromosome system and their thermal reaction norm (TRN). The latter varies from FM pattern (low and high temperatures producing females and males, respectively) to the opposite MF pattern through the FMF pattern (where males are produced at intermediate temperatures and females at the extremes). Asymmetrical sex reversal predicts, as detailed in Fig. 1, that under FM TRN pattern we should find ZW/ZZ systems in warmer environments than XX/XY systems, and the opposite difference is expected under MF pattern. For the FMF TRN pattern, where temperature has non-linear effect on sex and thus on the expected outcomes, whether and how the two GSD systems should differ in climatic distribution is predicted to depend on the extent of directional climate change each species has experienced (Fig. 1).

**Fig. 1.**
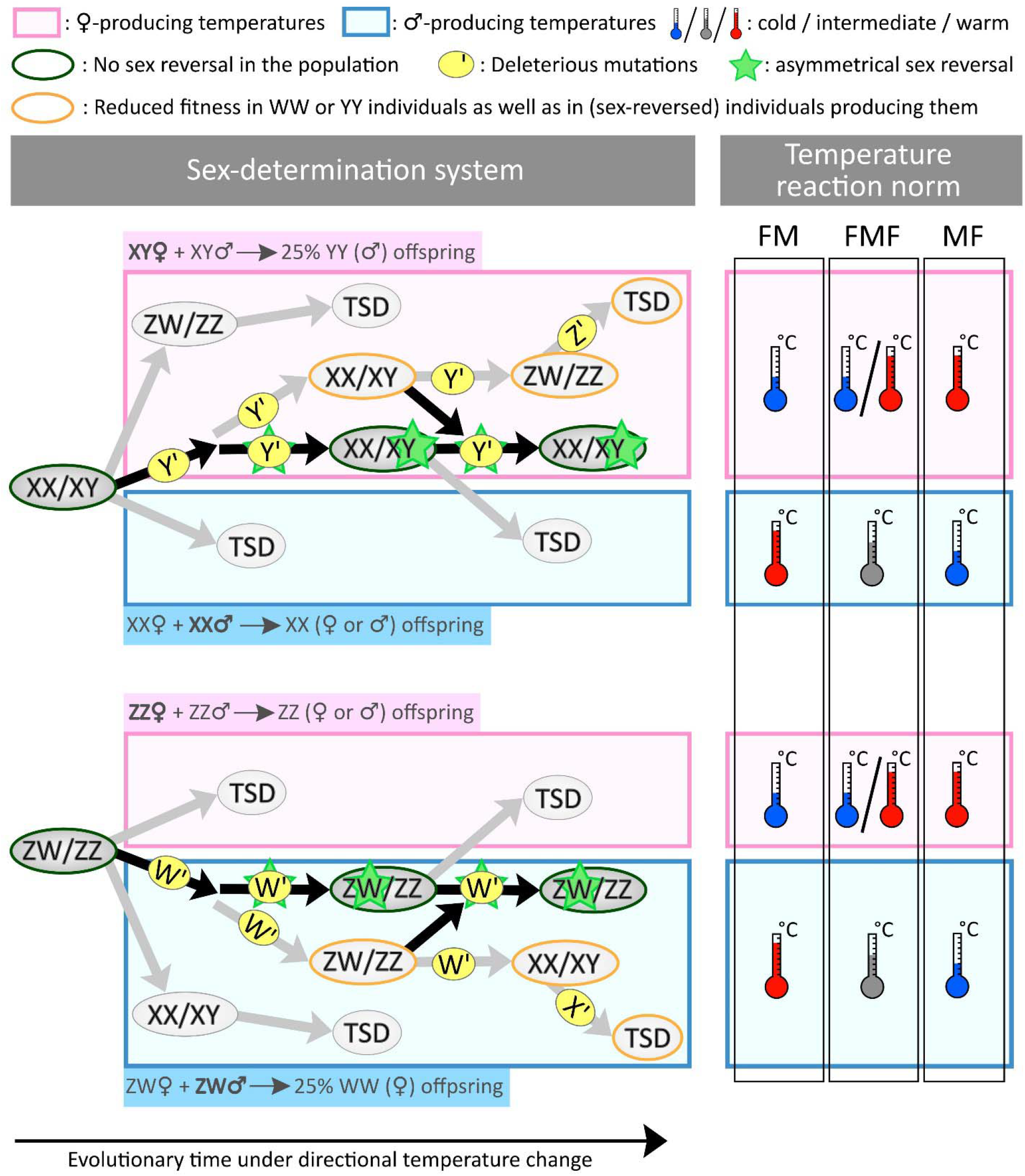
Hypothesized evolutionary pathways of sex-determination systems with thermal sex reversal. Directional climate change results in environmental temperatures that trigger male-to-female or female-to-male sex reversal (pink and light blue rectangles, respectively) depending on the pattern of thermal reaction norm (TRN) of the species. Changes in sex-chromosome frequencies cause turnovers between GSD systems, that become TSD when one sex chromosome disappears from the population. The paths denoted by grey arrows represent evolutionary scenarios predicted by earlier models (sex-ratio selection is key; ^13,15,25^. In systems denoted by orange circles, the population suffers increasing fitness reduction, and turnovers between these systems are only possible if the YY or WW genotype is viable and at least somewhat fertile; otherwise, early extinction is expected ^25^. The paths denoted by black arrows illustrate how asymmetrical sex reversal ^30^ may stabilize one heterogametic system, depending on the experienced environmental temperatures (symbolized by the thermometer icons on the right) and TRN. For example, if the TRN follows FM pattern, then high temperatures induce female-to-male sex reversal. In this scenario, ZW/ZZ systems are expected to be stabilized by sex-reversal resistance of the ZW genotype if deleterious mutations are accumulated on W; while in an XX/XY system, sex-reversed XX males will produce only XX offspring (benefitting from female-biased offspring sex ratio in a male-biased population), leading to the loss of Y. Therefore, under FM pattern, XX/XY systems may only persist in relatively colder habitats, while in warmer habitats ZW/ZZ systems should more likely occur. For FMF TRN pattern, predictions of climatic differences between GSD types depend on the extent of directional climate change. As long as these species experience moderate temperature increase, the climatic distribution of different GSD types may correspond with predictions for the FM pattern, but it should increasingly resemble the opposite MF pattern as warming progresses in their habitats. Although these evolutionary pathways may be altered by evolution of new TRNs ^14,16^, new sex-determination genes ^13^, and new sex chromosomes ^33–35^, for simplicity the illustrated scenarios assume no such changes.

In this study, we tested these predictions utilizing HerpSetDet, the most complete dataset of GSD types and TRN patterns across amphibians and reptiles ^1^, combining it with 16 variables calculated from temperature data of the WorldClim database ^32^ to describe the climate (thermal environment) of each species’ entire geographical distribution (Fig. 2). First, we used phylogenetic least-squares (PGLS) models to analyse how each climatic variable is related to the interaction between GSD type and TRN pattern. Second, we collected data on the temperatures likely experienced over the species’ geographical range during the period of sex determination. This was necessary because the sensitive window of sexual development, when environmental conditions may cause sex reversal, occurs during early ontogeny, but the timing of this period in nature is not known for many species. Thus, in order to pinpoint the PGLS models (run with the 16 climatic variables that were available for all species) that provide the most relevant test of our predictions, we used a smaller subset of species from both taxa (amphibians and reptiles) to identify the climatic variables (out of the 16) that correlate most strongly with the temperature of the sex-determination period.

**Fig. 2.**
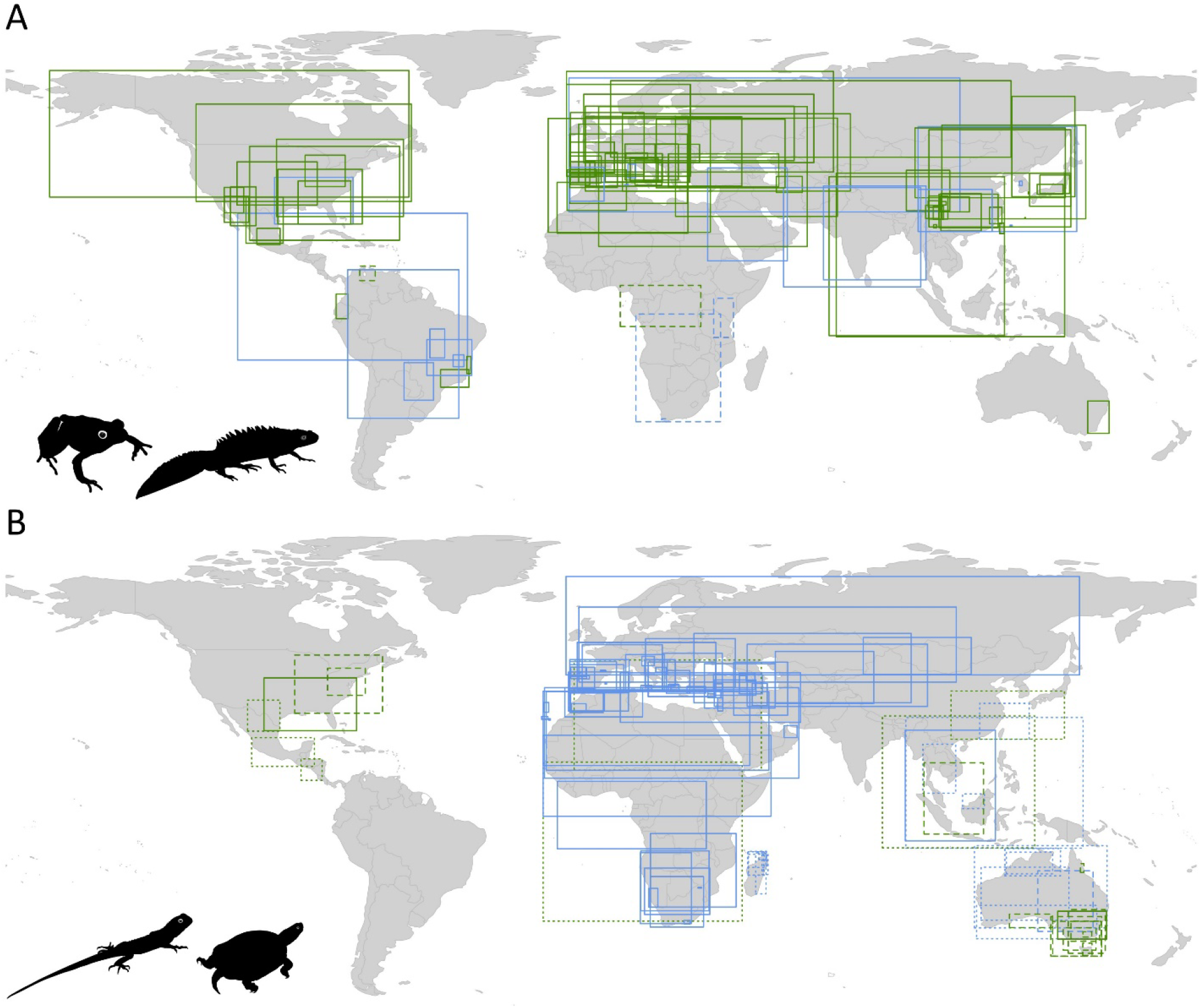
Global distribution of amphibian (A) and reptile (B) species included in the analyses. Rectangles represent the extent of the native distribution range of each species, calculated as described in the Methods section. Rectangle colours are denoting GSD type (green: XX/XY and blue: ZW/ZZ), while the border types are denoting TRN patterns (solid line: FM, dashed line: MF, dotted line: FMF). Note that the distribution ranges of 3 reptile species (*Emoia nigra, Hemidactylus frenatus, Lepidodactylus lugubris*) are not shown, because they occur (at least in part) on islands across the Pacific Ocean; therefore, the extent of their distribution range could not be denoted by one rectangle on the conventionally centred world map. The map was drawn by the ‘mapCountryData’ function of the ‘rworldmap’ R package, and rectangles were added by the ‘rect’ function in R 4.3.2. for each taxonomic class; these final maps were subsequently arranged on the figure panels in Inkscape 1.2.

## Results

Literature data enabled the assignment of TRN patterns to 208 species with known GSD type across the phylogeny of herpetofauna (Fig. 3A) distributed across all continents excepting Antarctica (Fig. 2.). In amphibians, six out of 16 models showed a significant interaction between GSD and TRN (Supplementary Table 1). In nine out of 16 models, species with the FM reaction norm occupied significantly warmer areas if they had the ZZ/ZW system compared to the XX/XY system, as predicted (Fig. 4; see details in Supplementary Table 1), while two models showed the predicted opposite difference between GSD systems among species with the MF reaction norm (Fig. 4; Supplementary Table 1). The climatic variable best correlating with the temperature of the sex-determining period across amphibians was the annual mean temperature in the warmest part of the distribution range (Supplementary Table 2); and this variable supported the predicted climatic differences between GSD types for FM species (Fig. 3B). The estimated temperatures of the sex-determining period had relatively strong correlations (i.e. stronger than the average across the 16 variables in amphibians) with most (six out of nine) climatic variables that showed the difference predicted for the FM pattern, and with both variables that supported our predictions for the MF pattern (Fig. 4; Supplementary Table 2).

**Fig. 3.**
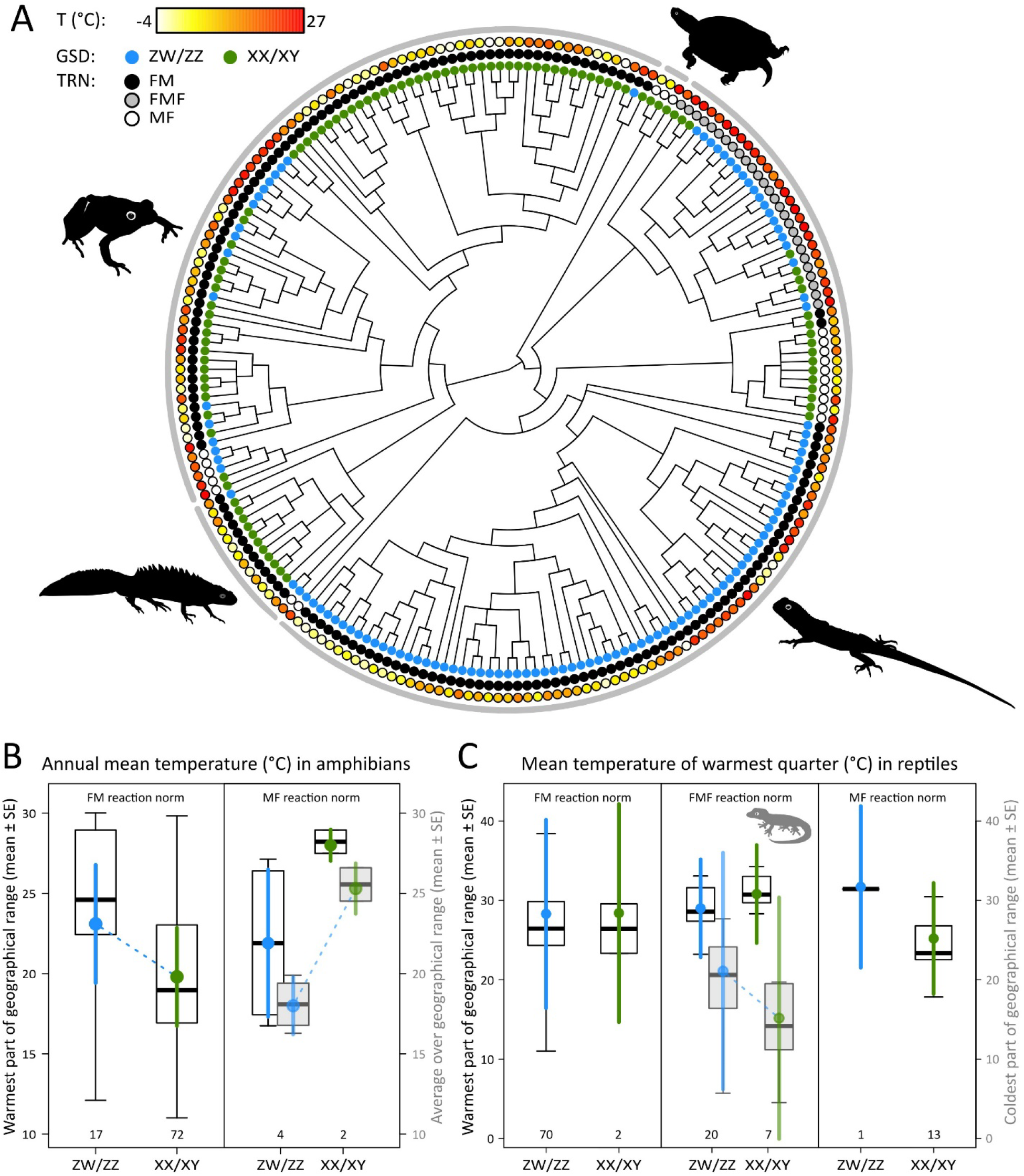
Phylogenetic relationships between climatic distribution, genotypic sex determination (GSD), and thermal reaction norm (TRN) in herpetofauna. Across the phylogeny (A), climate for each species is illustrated as the average annual mean temperature (T) over the geographical distribution range. The variable that correlates best with the temperature of the sex-determination period (black boxes with white fill) is the annual mean temperature in the warmest part of the distribution range for amphibians (B), and the mean temperature of the warmest quarter of the year in the warmest part of the distribution range for reptiles (C). Differences that support the theory of asymmetrical sex reversal are highlighted by dotted lines (P < 0.048) and illustrated by the third best-correlating proxy for the sex-determination period in each class (dark grey boxes with light grey fill): the annual mean temperature averaged over the distribution range for amphibians (B), and the mean temperature of the warmest quarter of the year in the coldest part of the distribution range for reptiles (C). In each box plot (B, C), the thick middle line, box, and whiskers represent the median, interquartile range, and data range, respectively; the coloured dots with error bars depict the mean and ± standard error estimated from phylogenetic least-squares models, and the number below each plot is the number of species for which sex-determination data are available. Figure panels were arranged and artwork was added in Inkscape 1.2.

**Fig. 4.**
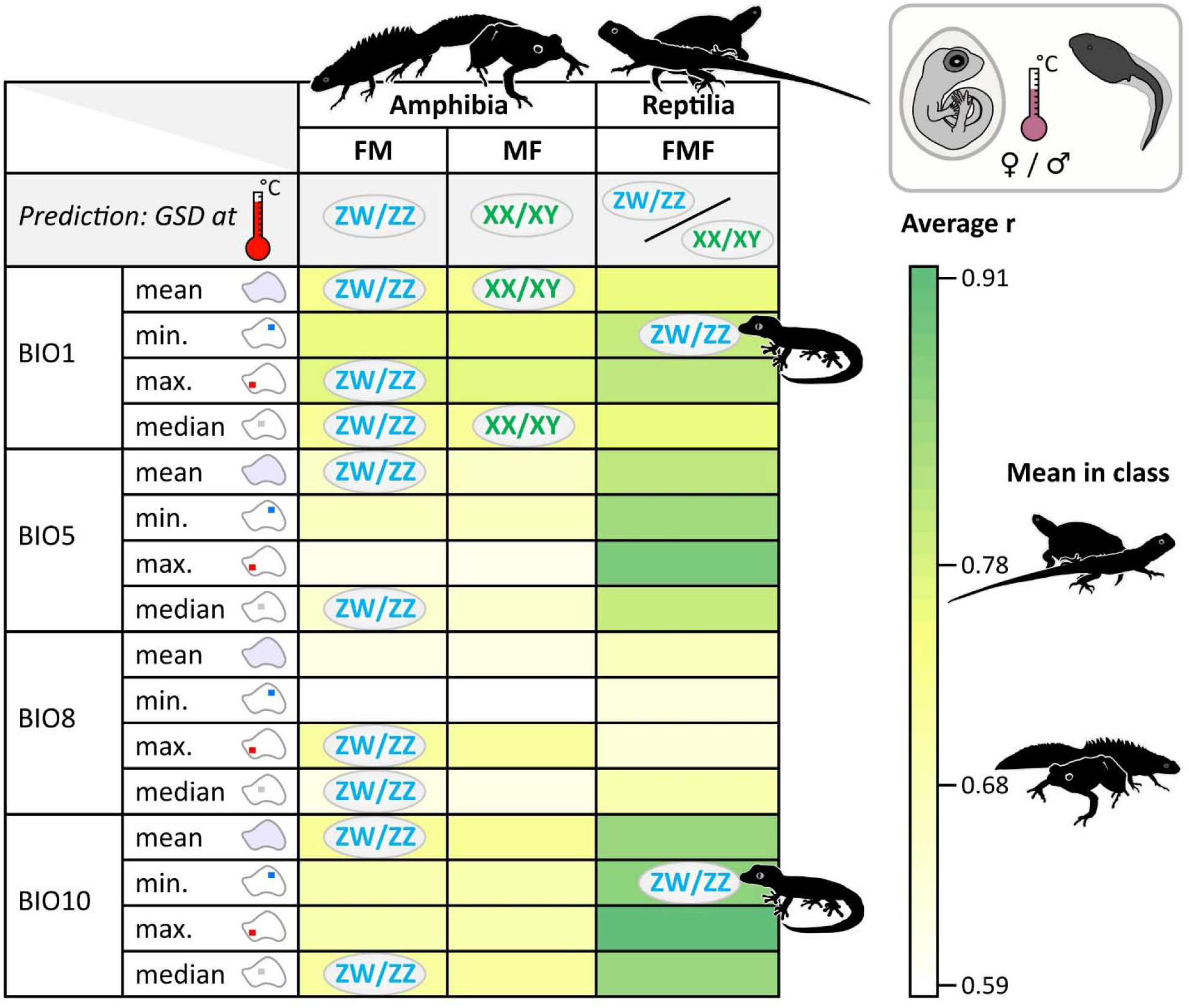
Overview of the results. of the 16 models and their estimated relevance for environmental temperatures around the sex-determining period. Grey ovals represent the sex-chromosome system that occurs at warmer temperatures according to each model for each TRN pattern. Polygons for each WorldClim variable illustrate how we calculated the mean, minimum (i.e. coldest area), maximum (i.e. warmest area), and median values over the native distribution range for annual mean temperature (BIO1), maximum temperature of the warmest month (BIO5), mean temperature of the wettest quarter (BIO8), and mean temperature of the warmest quarter (BIO10). Cell background colours indicate the correlation coefficients (Pearson’s r) averaged across corresponding SexClim variables (i.e. temperature variables calculated for the approximate sex-determining period for a subset of species) for each WorldClim variable in each taxonomic class.

In reptiles, the interaction between GSD and TRN was not significant in any analysis (Supplementary Table 1). Unexpectedly, in two out of 16 models, the coldest area of the distribution range was significantly warmer among species with the FMF reaction norm if they had the ZZ/ZW system compared to the XX/XY system (Fig. 3C, Fig. 4), while there were no differences between the two GSD systems among species with either FM or MF reaction norms (Fig. 3C, Supplementary Table 1). The climatic variable best correlating with the temperature of the sex-determining period across reptiles was the mean temperature of the warmest quarter of the year in the warmest part of the distribution range (Supplementary Table 2); however, the latter variable did not support any of the predicted effects of GSD and TRN (Supplementary Table 1, Fig. 4; Fig. 3C). The two climatic variables that showed a difference between GSD types within the FMF pattern had relatively strong correlations (i.e. stronger than the average across the 16 variables in reptiles) with the estimated temperatures of the sex-determining period (Fig. 4; Supplementary Table 2).

## Discussion

Our results demonstrate that the climatic distribution of ectothermic amniote species can be explained by the interplay between genotypic sex determination and clade-specific temperature reaction norm of sexual development, and for amphibians the observed differences conform to the predictions derived from the theory of asymmetrical sex reversal. These findings have several implications. First, they indicate that temperature-induced sex reversal may be an important driver of phylogeography and evolution of sex-determination systems across a wide range of species. Second, as contemporary climate change proceeds, the importance of temperature-induced sex reversal may increase, and the types of both GSD and TRN may determine which species would be most affected by its consequences, including climatic range shifts and even population extinctions. Finally, the different results we found in amphibians versus reptiles may either reflect true biological differences or may be simply due to variation in available sample size and thus statistical power. In either case, our study highlights the pressing need for collecting empirical data on GSD and especially TRN in many more species from a broad taxonomical spectrum, for a better understanding of both past and future evolutionary processes and conservation priorities.

In accordance with our predictions, we found that amphibian species with ZW/ZZ systems occupied significantly warmer areas than those with XX/XY systems in clades with FM TRN pattern, while the opposite pattern was found in clades with MF pattern. These differences were found with several climatic variables, which were not a random subset of the variables we investigated: mostly they showed higher than average correlation with temperature variables measured within the period of larval development in amphibians. These differences not only follow the predictions of asymmetrical sex reversal, but to our knowledge, no other current theory explains them. In contrast, we found none of the predicted differences in reptiles with FM or MF pattern. The simplest explanation for these contrasting results may lie in the heterogeneity of sample sizes available from the literature. The comparison most persistently supported by our analyses was the one for which we had the highest power (i.e. N > 15 species for each GSD system): within FM amphibians. For reptiles, all three comparisons had limited power because the number of species was very low (≤ 7) for one of the two GSD systems within each comparison. However, an alternative explanation is that reptiles may truly be less conforming to asymmetrical sex reversal compared to amphibians. Potential reasons for this include maternal thermoregulation (e.g. basking ^36^) and nest-site choice ^37^ in viviparous and oviparous reptiles, respectively), which may allow for higher control over the thermal environment of their offspring compared to amphibians where free-moving aquatic larvae are most typical ^38^; however, aquatic larvae may also express stage-dependent temperature preferences within the limitations of their microhabitat ^39,40^. To confront these alternatives and to test the generality of the asymmetrical sex-reversal theory, we urge empirical work especially on species in clades where GSD occurs and the effects of temperature on sexual development have not yet been tested. Because temperature-induced sex reversal can occur not only in herpetofauna but also in other taxa like fish ^22,41,42^, studies of marine and freshwater temperatures across various water depths could further clarify the role of temperature-induced sex reversal in the phylogeography and evolution of GSD species.

Unexpectedly, we found in reptiles with FMF TRN pattern that, during the warmest quarter of the year (the period with highest overall correlation with environmental temperatures during embryonic development in reptiles; see Fig. 4), the coldest parts of the geographical distribution ranges of ZW/ZZ species were significantly warmer compared to those of XX/XY species. This difference resembles the pattern expected under FM reaction norm, and thus it might suggest that environmental temperatures of the studied species in clades with FMF are not yet so high that would induce male-to-female sex reversal, and instead, may currently experience temperatures around the lower threshold (i.e. female-producing temperatures below and male-producing temperatures above this threshold). In our dataset, the FMF TRN pattern was only assigned to geckos (families Eublepharidae and Gekkonidae), due to the lack of GSD data in other FMF species or to the ambiguity of TRN patterns in other families where FMF occurs. We had data on the timing of early embryonic development for some of these species, and the mean temperature across their natural distribution range during this period (17-27°C) was indeed closer to the lower temperature threshold (the lower and higher TRN thresholds in these families are around 26-30°C and 32-34°C, facilitating the development of phenotypic females below, males in between and females above, respectively ^1^), conforming to the above suggestion. It is therefore possible that temperature-induced (male-to-female) sex reversal might predominantly occur at the coldest parts of the geographical distribution ranges in the studied gecko species, contributing to the temperature difference between species with XX/XY and ZW/ZZ systems found here. However, empirical data are currently missing to verify this explanation.

Our results highlight the importance of collecting data on the occurrence of sex chromosomes in parallel with data on the effects of temperature on sexual development in ectothermic vertebrate species. Empirical studies so far, especially in reptiles, predominantly have been focusing on either genotypic or temperature-dependent sex determination, and once a species was categorized as either GSD or TSD, researchers rarely attempted to evaluate if sex chromosomes and environmental temperatures could both influence sexual development; although such rare attempts revealed cryptic patterns ^28,43–45^. Moreover, various studies assessing TSD applied only a limited number of incubation temperatures and made conclusions on limited sample sizes (see “TSD_note” in HerpSexDet ^1^); and it can be difficult to infer if sex-ratio bias was caused by sex reversal or sex-biased mortality (see common methodological constrains in ^30^). We suspect that these limitations are largely responsible for the significant lack of fundamental empirical information regarding the potential temperature-sensitivity of sexual development under GSD. Besides conducting laboratory experiments, it is important to uncover sex-reversal frequencies in wild populations as well, by comparing the genotypic sex with the phenotypic sex in each captured individual (^17,28,44^; see further reports in ^1^); and such studies should account for the possibility that certain chemical pollutants may also induce sex reversal (see examples in ^30^). Collecting such empirical data is crucial for understanding how and to what extent environmental conditions influence the evolution of sex determination and its consequences for demography and population persistence.

While previous empirical studies suggested that GSD and TSD systems in reptiles are associated with different climatic conditions ^5,43^, here we demonstrated for the first time that within GSD, even XX/XY and ZW/ZZ systems can be associated with different environmental temperatures. Our results on the climatic distribution of XX/XY and ZW/ZZ systems suggest that directional temperature changes (either via climate change, geographical range shifts or both) may influence transitions between different sex-determination systems in a non-random way, favouring one sex-chromosome system over the other, depending on the shape of the TRN featured by the given taxon. Evaluating if species are able to adapt fast enough to currently accelerating climatic changes is one of the most pressing biodiversity-conservation problems. We propose that taking asymmetrical sex reversal into account can help to pinpoint the taxa most vulnerable to rising environmental temperatures.

## Methods

### Sex determination

We collected species-specific data on the type of GSD and TRN of sexual development from the HerpSexDet database version 1.1. ^1^. For the sake of the analyses, we classified both simple (i.e. one sex chromosome pair) and complex (i.e. multiple sex chromosomes) male-heterogametic systems as “XX/XY”, and simple and complex female-heterogametic systems as “ZW/ZZ”, and excluded species with rarer GSD types. Because TRN data were not available for most GSD species, we assigned one of the three common TRNs (i.e. FM, FMF or MF, corresponding to the sex predominantly produced at different temperature ranges from low to high: F for females and M for males) to each taxonomic family, provided that one TRN pattern was reported for at least two species (regardless of GSD being identified in them or not) and no other TRN pattern was reported in that family. When more than one TRN pattern occurred in a family, we aimed to find phylogenetic clades below the family level that conformed to the above criteria (according to the Open Tree of Life Synthetic Tree ^46^). When TRN information was available for only one species in the family, we assigned that TRN pattern to that family only if it was not conflicted by TRN data in the phylogenetically closest family. Thus, we assigned the species-specific TRN for each species where it was available, and the TRN characteristic to its clade to each species with unknown TRN (see detailed justification in Supplementary Table 3).

### Spatial distribution

We collected data on the species’ distribution ranges, preferably as spatial polygons obtained from the IUCN Red List of Threatened Species (spatial database version 2022-2 ^47^; hereafter IUCN). Some species were not present in the IUCN datasets or we judged that the available data were insufficient for further processing: the polygons were too small for masking the raster grid cells (see the “Climatic variables” section below), or the coordinates assigned to the polygons were noticed to be inaccurate. In these cases, if available, spatial occurrence data were obtained from the database of the Global Biodiversity Information Facility (hereafter GBIF ^48^). We focused on the native distribution range of species where the species are currently extant and not introduced. Specifically, IUCN spatial polygons categorized as ‘introduced’, ‘assisted colonisation’ or ‘vagrant’ were excluded and the seasonality was required to be set either as ‘resident’ or ‘breeding season’; and only GBIF datapoints categorized as ‘present’ with coordinate uncertainties below 10 km were included. Because GBIF provides no information on whether a population is introduced or not, we checked for introduced populations of the concerned reptile species in The Reptile Database ^49^, but did not find indication of introduced populations of the species with GBIF data, so we assumed that all datapoints represented native occurrences (we did not find such information for the concerned amphibian species elsewhere). We excluded those species from further analyses that had less than 10 GBIF datapoints after this filtering. All in all, spatial distribution data were collected from IUCN for 93 amphibian and 107 reptile species, and from GBIF for two amphibian and six reptile species. Geographical occurrence shapes were defined either as polygons provided by IUCN, or as GBIF datapoints converted to shape files using the ‘writeOGR’ function of the ‘rgdal’ package ^50^.

### Climatic variables

Data calculations from spatial datasets were performed in R 4.2.1 ^51^. We obtained temperature data across a 30-year period (1970-2000) in five-minute spatial resolution from the WorldClim database version 2.1. ^32^. Out of the available 19 “bioclimatic” variables, we used those that we deemed likely to encompass temperature data relevant to the offspring-development season of herpetofauna: annual mean temperature (BIO1), maximum temperature of the warmest month (BIO5), mean temperature of the wettest quarter (BIO8), and mean temperature of the warmest quarter (BIO10). We cut out the geographical occurrence shapes from each raster band using the ‘mask’ function (‘raster’ package ^52^). From these cuts, we obtained the temperature values with the ‘getValues’ function (‘raster’ package ^52^) and subsequently calculated the mean, minimum, maximum and median values of each of the 4 WorldClim rasters (hereafter these 16 variables will be referred to as WorldClim variables) across each species’ native distribution range. For visualisation purposes, the westernmost, easternmost, southernmost and northernmost coordinates of the distribution were calculated by the ‘extent’ function of the ‘raster’ package ^52^ for polygons downloaded from the IUCN database; while for GBIF datapoints, they were calculated as the minimum and maximum values of the X and Y coordinates across datapoints, respectively (based on these calculations, spatial distribution of the studied species is shown on Fig. 2).

The period when sexual development is sensitive to environmental temperatures (referred to as “sex-determining period” across the main text) occurs during early ontogeny, during the larval phase in amphibians ^53–55^ and the embryonic phase in reptiles ^28,56,57^. Temperature during this period would be the most relevant to our study; however, the timing of this period is unknown for many species. Therefore, we collected information from the literature on the timing of the larval/embryonic season (in months) for a subset of the analysed species. To approximate the sex-determining period, we estimated the timing of the larval/embryonic period in two ways. First, we took the month(s) when mating occurs in species with external fertilization, and the month(s) when egg-laying occurs in oviparous species. For viviparous species (where sex determination occurs before birth), we took the month(s) when fertilization occurs or gravid females appear, or the mating season if the previous two were not reported. Second, we extended the period described in the previous two sentences by one month, because the length of larval/embryonic developmental phase, the timing of the temperature-sensitive window within that phase, and the developmental stage of embryos at egg laying can differ between species. For both approximations of the sex-determination period (i.e. the non-extended and the extended larval/embryonic periods), we used monthly average temperature data in five-minute resolution from WorldClim database version 2.1, ^32^ to calculate mean, median, minimum and maximum temperatures across the native range (hereafter SexClim variables) for each month of the sex-determining period, by the same computational methods as described above for the WorldClim variables. Subsequently, we calculated the mean and median temperatures for the entire sex-determining period (i.e. mean of the monthly mean temperatures across the native range, and median of the monthly median temperatures across the native range, respectively) as well as for the coldest and the warmest month within that period. Furthermore, we assessed potential temperature extremes for larval/embryonic development by calculating the average temperatures of the coldest and warmest parts of the natural distribution range during the coldest and warmest month, respectively. In those species where the embryonic/larval season reportedly differed along a South-North cline (10 amphibian species), we divided the natural distribution range into maximum three subranges at equal distances along its longitudinal extent using the ‘gIntersection’ function (‘rgeos’ package ^58^), and calculated the related SexClim variables separately for each subrange. Then, we calculated the relevant statistics (mean, median, minimum or maximum, depending on the variable) from these subranges and assigned a single value for each SexClim variable for each concerned species.

We excluded the cave-breeding amphibian genus Hydromantes ^59^ as well as all viviparous sea snakes from our dataset, because the land-surface temperatures of the WorldClim database might not be informative for conditions experienced by the offspring of species inhabiting caves or relatively deep sea water. Although many amphibian species have aquatic larvae, those almost exclusively occupy shallow freshwater bodies, where water temperatures correlate well with air temperatures ^60,61^. Therefore, we assumed that the WorldClim data adequately reflect the relevant climatic differences not only among reptiles (whose offspring typically develop in terrestrial nests) but also among amphibian species. Our final dataset, for which we obtained WorldClim variables, consisted of 95 amphibian and 113 reptile species with either male or female heterogamety as well as either species-or clade-specific information on the TRN of sexual development. Out of these, we found SexClim data for 74 amphibian and 47 reptile species. The final dataset is available upon request.

### Statistical analyses

All statistical analyses were run in R 4.3.2 ^62^. We built phylogenetic generalized least squares (PGLS) models ^63^ using the ‘gls’ function of the ‘nlme’ package ^64^. To control for the phylogenetic relatedness among taxa, we used the Open Tree of Life Synthetic Tree ^46^ that we accessed using the ‘rotl’ package ^65^. The resulting topology is available upon request. Since composite phylogenies do not have true branch lengths, we used Grafen’s method ^66^ to generate branch lengths using the ‘ape’ package ^67^. In each model we used the phylogenetic signal (i.e. Pagel’s λ) as estimated by the maximum-likelihood method for that model. In each PGLS model, the dependent variable was one of the 16 WorldClim variables, and the fixed effects were the type of GSD (XX/XY or ZZ/ZW), the type of TRN (MF, FMF, or FMF), and their interaction. Because the FMF reaction norm is not known to occur in any amphibian families, we analysed amphibians and reptiles separately. Due to differences in sample size, we allowed for variances to differ between groups (among the four combinations of GSD and TRN in amphibians, and among the three TRN types in reptiles; we could not estimate variances for each combination of GSD and TRN in reptiles because one of these combinations had no variance, i.e. N=1). To test our predictions, we used pre-planned comparisons and report the effect sizes with 95% confidence intervals (corresponding to a significance level of 5%) as recommended for evolutionary-ecological studies with multiple comparisons^68,69^. From each model, we estimated the difference (linear contrast) between XX/XY and ZZ/ZW systems within each type of TRN using the ‘emmeans’ function of the ‘emmeans’ package ^70^. We present the detailed results of all models in Supplementary Table 1.

We analysed the WorldClim variables to maximise sample size, as these data were available for all species in our dataset (N = 208) while we had SexClim data for a smaller subset (N = 120). To assess how well each WorldClim variable may reflect the variation across species in the temperatures of the sex-determining period, we tested the pairwise Pearson correlation between each of the 16 WorldClim variables and each of those SexClim variables that referred to the same aspect of climatic distribution (e.g. mean, median, etc.) separately for amphibians and reptiles (Supplementary Table 2).

## Data availability

The dataset and and the phylogenetic tree used in this study will be published after acceptance of the related article. Until then, those are available from the authors upon reasonable request.

## Acknowledgement

We thank Mihály B. Rusz for his help in data collection. This research was funded in part by the Austrian Science Fund (FWF) [grant ESP 239-B]. For open access purposes, the author has applied a CC BY public copyright license to any author-accepted manuscript version arising from this submission. V.B. was supported by the National Research, Development and Innovation Office of Hungary (K-135016).

## Contributions

N.E. conceived the concept of the study, carried out data collection, calculated the climatic variables and wrote the first draft of the manuscript. V. B. fine-tuned the study goals, carried out the statistical analyses and revised the manuscript.

