## Supplementary Table 1 for "Interplay of genotypic and thermal sex determination shapes climatic distribution in herpetofauna"

```
Supplementary Table 1. Results of phylogenetic least squares (PGLS) models for each climatic variable. In each model, the intercept refers to ZZ/ZW system and FM pattern; the phylogenetic signal (λ) is estimated by maximum likelihood. Within each TRN pattern, for each type of GSD and also for the difference between the two GSD types, model-estimated means are reported with standard errors (SE), Satterthwaite degrees of freedom (df), and 95% lower and upper confidence limits (CL).

Annual average temperature (BIO1) mean over geographical range

Amphibians:

λ = 0.78

PGLS coefficients:
                estimate        SE   t-value p-value
Intercept      17.059455  5.285546  3.227567  0.0017
 - XX/XY       -4.093992  1.681146 -2.435240  0.0168
 - MF           0.935104  4.304267  0.217250  0.8285
 - XX/XY × MF  11.373369  2.015891  5.641858  0.0000

Estimated group means:

TRN = FM:
 GSD     mean   SE   df lower CL upper CL
 ZZ/ZW   17.1 5.29 3.73     1.95     32.2
 XX/XY   13.0 5.26 2.76    -4.61     30.5

TRN = MF:
 GSD     mean   SE   df lower CL upper CL
 ZZ/ZW   18.0 1.79 1.92     9.99     26.0
 XX/XY   25.3 1.57 1.58    16.49     34.1

Pairwise comparisons:

TRN = FM:
 contrast          estimate   SE    df t-value p-value  lower CL upper CL
 (ZZ/ZW) - (XX/XY)     4.09 1.68 52.89   2.435  0.0183     0.722     7.47

TRN = MF:
 contrast          estimate   SE    df t-value p-value  lower CL upper CL
 (ZZ/ZW) - (XX/XY)    -7.28 1.10  5.59  -6.645  0.0007   -10.008    -4.55


Reptiles:

λ = 0.75

PGLS coefficients:
                 estimate        SE    t-value p-value
Intercept       15.083691  7.717290  1.9545320  0.0532
 - XX/XY        -0.115125  9.816143 -0.0117281  0.9907
 - FMF           9.323233  9.093514  1.0252618  0.3076
 - MF            6.438407 12.146176  0.5300769  0.5972
 - XX/XY × FMF  -3.052826  9.938691 -0.3071658  0.7593
 - XX/XY × MF   -6.163488 12.895673 -0.4779501  0.6337

Estimated group means:

TRN = FM:
 GSD     mean    SE   df lower CL upper CL
 ZZ/ZW   15.1  7.72 2.69  -11.128     41.3
 XX/XY   15.0  6.99 5.30   -2.699     32.6

TRN = FMF:
 GSD     mean    SE   df lower CL upper CL
 ZZ/ZW   24.4  5.22 2.09    2.878     45.9
 XX/XY   21.2  5.28 2.21    0.465     42.0

TRN = MF:
 GSD     mean    SE   df lower CL upper CL
 ZZ/ZW   21.5 10.93 2.93  -13.734     56.8
 XX/XY   15.2  6.94 2.01  -14.417     44.9

Pairwise comparisons:

TRN = FM:
 contrast          estimate    SE    df t-value p-value  lower CL upper CL
 (ZZ/ZW) - (XX/XY)    0.115  9.82  3.64   0.012  0.9913    -28.2274     28.5

TRN = FMF:
 contrast          estimate    SE    df t-value p-value  lower CL upper CL
 (ZZ/ZW) - (XX/XY)    3.168  1.56 21.27   2.036  0.0544      -0.0652      6.4

TRN = MF:
 contrast          estimate    SE    df t-value p-value  lower CL upper CL
 (ZZ/ZW) - (XX/XY)    6.279 12.33  2.71   0.509  0.6492  -35.4653     48.0

Annual average temperature (BIO1) minimum over geographical range

Amphibians:

λ = 0.85

PGLS coefficients:
                estimate        SE    t-value p-value
Intercept       0.892112  9.115914  0.0978632  0.9223
 - XX/XY       -1.856201  2.465502 -0.7528691  0.4535
 - MF          11.945974  6.171615  1.9356316  0.0560
 - XX/XY × MF   3.445420  6.241338  0.5520322  0.5823

Estimated group means:

TRN = FM:
 GSD     mean   SE   df lower CL upper CL
 ZZ/ZW  0.892 9.12 7.16   -20.57     22.4
 XX/XY -0.964 8.49 5.73   -21.99     20.1

TRN = MF:
 GSD     mean   SE   df lower CL upper CL
 ZZ/ZW 12.838 7.20 2.75   -11.30     37.0
 XX/XY 14.427 1.29 1.82     8.32     20.5

Pairwise comparisons:

TRN = FM:
 contrast          estimate   SE    df t-value p-value  lower CL upper CL
 (ZZ/ZW) - (XX/XY)     1.86 2.47 10.98   0.753  0.4674     -3.57     7.28

TRN = MF:
 contrast          estimate   SE    df t-value p-value  lower CL upper CL
 (ZZ/ZW) - (XX/XY)    -1.59 6.15  2.24  -0.258  0.8181    -25.55    22.37


Reptiles:

λ = 0.90

PGLS coefficients:
                 estimate        SE    t-value p-value
Intercept        6.316676  17.51982  0.3605445  0.7192
 - XX/XY         0.241855  21.33869  0.0113341  0.9910
 - FMF          12.021151  24.28946  0.4949122  0.6217
 - MF            7.249348  21.26098  0.3409697  0.7338
 - XX/XY × FMF  -8.705016  21.62248 -0.4025910  0.6881
 - XX/XY × MF   -5.126908  22.30957 -0.2298076  0.8187

Estimated group means:

TRN = FM:
 GSD     mean   SE   df lower CL upper CL
 ZZ/ZW   6.32 17.5 71.5    -28.6     41.2
 XX/XY   6.56 14.4 71.5    -22.2     35.4

TRN = FMF:
 GSD     mean   SE   df lower CL upper CL
 ZZ/ZW  18.34 17.8 29.4    -18.0     54.6
 XX/XY   9.87 17.8 29.4    -26.5     46.3

TRN = MF:
 GSD     mean   SE   df lower CL upper CL
 ZZ/ZW  13.57 16.1 15.5    -20.8     47.9
 XX/XY   8.68 11.0 15.5    -14.7     32.0

Pairwise comparisons:

TRN = FM:
 contrast          estimate    SE   df t-value p-value  lower CL upper CL
 (ZZ/ZW) - (XX/XY)   -0.242 21.34 71.5  -0.011  0.9910    -42.78     42.3

TRN = FMF:
 contrast          estimate    SE   df t-value p-value  lower CL upper CL
 (ZZ/ZW) - (XX/XY)    8.463  3.49 29.4   2.424  0.0217      1.33     15.6

TRN = MF:
 contrast          estimate    SE   df t-value p-value  lower CL upper CL
 (ZZ/ZW) - (XX/XY)    4.885 18.49 15.5   0.264  0.7951    -34.41     44.2

Annual average temperature (BIO1) maximum over geographical range

Amphibians:

λ = 0.69

PGLS coefficients:
                estimate        SE   t-value p-value
Intercept      23.136806  3.683397  6.281377  0.0000
 - XX/XY       -3.324759  1.407811 -2.361651  0.0203
 - MF          -1.205956  3.835190 -0.314445  0.7539
 - XX/XY × MF   9.391214  3.881892  2.419236  0.0175

Estimated group means:

TRN = FM:
 GSD     mean    SE   df lower CL upper CL
 ZZ/ZW   23.1 3.683 5.99    14.12     32.2
 XX/XY   19.8 3.063 6.86    12.54     27.1

TRN = MF:
 GSD     mean    SE   df lower CL upper CL
 ZZ/ZW   21.9 4.588 3.21     7.85     36.0
 XX/XY   28.0 0.975 1.93    23.65     32.3

Pairwise comparisons:

TRN = FM:
 contrast          estimate   SE   df t-value p-value  lower CL upper CL
 (ZZ/ZW) - (XX/XY)     3.32 1.41 7.96   2.362  0.0460    0.0755     6.57

TRN = MF:
 contrast          estimate   SE   df t-value p-value  lower CL upper CL
 (ZZ/ZW) - (XX/XY)    -6.07 3.93 2.65  -1.545  0.2315  -19.5355     7.40


Reptiles:

λ = 0.88

PGLS coefficients:
                 estimate        SE    t-value p-value
Intercept       21.173349  9.912907  2.1359374  0.0350
 - XX/XY         1.254249 12.136567  0.1033446  0.9179
 - FMF           5.639778 11.265768  0.5006119  0.6177
 - MF            5.460697 14.820550  0.3684544  0.7133
 - XX/XY × FMF  -1.250098 12.200819 -0.1024602  0.9186
 - XX/XY × MF   -8.174033 15.420368 -0.5300803  0.5972

Estimated group means:

TRN = FM:
 GSD     mean    SE    df lower CL upper CL
 ZZ/ZW   21.2  9.91  7.86    -1.76     44.1
 XX/XY   22.4  8.27 10.55     4.13     40.7

TRN = FMF:
 GSD     mean    SE    df lower CL upper CL
 ZZ/ZW   26.8  5.89  6.18    12.50     41.1
 XX/XY   26.8  5.91  6.29    12.51     41.1

TRN = MF:
 GSD     mean    SE    df lower CL upper CL
 ZZ/ZW   26.6 13.20  5.30    -6.74     60.0
 XX/XY   19.7  8.90  4.60    -3.79     43.2

Pairwise comparisons:

TRN = FM:
 contrast          estimate    SE    df t-value p-value  lower CL upper CL
 (ZZ/ZW) - (XX/XY) -1.25425 12.14  8.91  -0.103  0.9200    -28.75    26.24

TRN = FMF:
 contrast          estimate    SE    df t-value p-value  lower CL upper CL
 (ZZ/ZW) - (XX/XY) -0.00415  1.25 28.19  -0.003  0.9974     -2.56     2.56

TRN = MF:
 contrast          estimate    SE    df t-value p-value  lower CL upper CL
 (ZZ/ZW) - (XX/XY)  6.91978 15.08  5.13   0.459  0.6652    -31.55    45.39

Annual average temperature (BIO1) median over geographical range

Amphibians:

λ = 0.78

PGLS coefficients:
                estimate        SE   t-value p-value
Intercept      17.249232  5.460514  3.158902  0.0021
 - XX/XY       -4.059751  1.747121 -2.323680  0.0224
 - MF           0.980880  4.385961  0.223641  0.8235
 - XX/XY × MF  11.545438  2.191778  5.267613  0.0000

Estimated group means:

TRN = FM:
 GSD     mean   SE   df lower CL upper CL
 ZZ/ZW   17.2 5.46 3.76     1.69     32.8
 XX/XY   13.2 5.50 2.77    -5.16     31.5

TRN = MF:
 GSD     mean   SE   df lower CL upper CL
 ZZ/ZW   18.2 2.00 1.91     9.23     27.2
 XX/XY   25.7 2.07 1.58    14.13     37.3

Pairwise comparisons:

TRN = FM:
 contrast          estimate   SE    df t-value p-value  lower CL upper CL
 (ZZ/ZW) - (XX/XY)     4.06 1.75 69.63   2.324  0.0231     0.575     7.54

TRN = MF:
 contrast          estimate   SE    df t-value p-value  lower CL upper CL
 (ZZ/ZW) - (XX/XY)    -7.49 1.31  5.01  -5.734  0.0022   -10.839    -4.13


Reptiles:

λ = 0.74

PGLS coefficients:
                 estimate        SE    t-value p-value
Intercept       15.255981  7.662777  1.9909206  0.0490
 - XX/XY        -0.216001  9.784316 -0.0220763  0.9824
 - FMF           9.362572  9.082510  1.0308353  0.3049
 - MF            6.453396 11.951157  0.5399809  0.5903
 - XX/XY × FMF  -2.895018  9.916245 -0.2919470  0.7709
 - XX/XY × MF   -5.975900 12.722481 -0.4697119  0.6395

Estimated group means:

TRN = FM:
 GSD     mean    SE   df lower CL upper CL
 ZZ/ZW   15.3  7.66 2.71  -10.696     41.2
 XX/XY   15.0  7.00 5.50   -2.464     32.5

TRN = FMF:
 GSD     mean    SE   df lower CL upper CL
 ZZ/ZW   24.6  5.29 2.10    2.917     46.3
 XX/XY   21.5  5.35 2.23    0.604     42.4

TRN = MF:
 GSD     mean    SE   df lower CL upper CL
 ZZ/ZW   21.7 10.70 3.02  -12.187     55.6
 XX/XY   15.5  6.75 2.04  -12.971     44.0

Pairwise comparisons:

TRN = FM:
 contrast          estimate    SE    df t-value p-value  lower CL upper CL
 (ZZ/ZW) - (XX/XY)    0.216  9.78  3.72   0.022  0.9835   -27.784    28.22

TRN = FMF:
 contrast          estimate    SE    df t-value p-value  lower CL upper CL
 (ZZ/ZW) - (XX/XY)    3.111  1.61 21.35   1.930  0.0670    -0.238     6.46

TRN = MF:
 contrast          estimate    SE    df t-value p-value  lower CL upper CL
 (ZZ/ZW) - (XX/XY)    6.192 12.06  2.79   0.514  0.6455   -33.896    46.28


Maximum temperature of warmest month (BIO5) mean over geographical range

Amphibians:

λ = 0.75

PGLS coefficients:
                estimate        SE   t-value p-value
Intercept      30.014485  3.058337  9.813990  0.0000
 - XX/XY       -3.293068  1.228432 -2.680708  0.0087
 - MF           0.104196  2.776989  0.037521  0.9702
 - XX/XY × MF   4.987291  3.302749  1.510042  0.1345

Estimated group means:

TRN = FM:
 GSD     mean   SE   df lower CL upper CL
 ZZ/ZW   30.0 3.06 5.09     22.2     37.8
 XX/XY   26.7 3.57 5.51     17.8     35.6

TRN = MF:
 GSD     mean   SE   df lower CL upper CL
 ZZ/ZW   30.1 3.43 2.88     18.9     41.3
 XX/XY   31.8 1.01 1.85     27.1     36.5

Pairwise comparisons:

TRN = FM:
 contrast          estimate   SE    df t-value p-value  lower CL upper CL
 (ZZ/ZW) - (XX/XY)     3.29 1.23 14.61   2.681  0.0174     0.669     5.92

TRN = MF:
 contrast          estimate   SE    df t-value p-value  lower CL upper CL
 (ZZ/ZW) - (XX/XY)    -1.69 2.73  2.19  -0.622  0.5927   -12.489     9.10


Reptiles:

λ = 0.88

PGLS coefficients:
                 estimate        SE   t-value p-value
Intercept       30.375509  8.449093  3.595121  0.0005
 - XX/XY         0.880883 10.335721  0.085227  0.9322
 - FMF           2.453713 10.206647  0.240403  0.8105
 - MF            6.202060 10.086054  0.614914  0.5399
 - XX/XY × FMF  -1.482807 10.416790 -0.142348  0.8871
 - XX/XY × MF   -9.655304 10.658937 -0.905841  0.3671


Estimated group means:

TRN = FM:
 GSD     mean   SE   df lower CL upper CL
 ZZ/ZW   30.4 8.45 5.99      9.7     51.1
 XX/XY   31.3 7.04 7.98     15.0     47.5

TRN = FMF:
 GSD     mean   SE   df lower CL upper CL
 ZZ/ZW   32.8 6.18 5.86     17.6     48.0
 XX/XY   32.2 6.20 5.96     17.0     47.4

TRN = MF:
 GSD     mean   SE   df lower CL upper CL
 ZZ/ZW   36.6 7.48 4.43     16.6     56.6
 XX/XY   27.8 5.05 3.82     13.5     42.1

Pairwise comparisons:

TRN = FM:
 contrast          estimate    SE    df t-value p-value  lower CL upper CL
 (ZZ/ZW) - (XX/XY)   -0.881 10.34  6.77  -0.085  0.9345    -25.49    23.73

TRN = FMF:
 contrast          estimate    SE    df t-value p-value  lower CL upper CL
 (ZZ/ZW) - (XX/XY)    0.602  1.30 29.70   0.464  0.6460     -2.05     3.25

TRN = MF:
 contrast          estimate    SE    df t-value p-value  lower CL upper CL
 (ZZ/ZW) - (XX/XY)    8.774  8.55  4.28   1.027  0.3591    -14.35    31.90


Maximum temperature of warmest month (BIO5) minimum over geographical range

Amphibians:

λ = 0.51

PGLS coefficients:
                estimate        SE    t-value p-value
Intercept      14.505300  4.776357  3.0368960  0.0031
 - XX/XY       -0.698692  2.286591 -0.3055604  0.7606
 - MF           7.951753  5.016641  1.5850751  0.1164
 - XX/XY × MF  -2.548661  5.122335 -0.4975584  0.6200

Estimated group means:

TRN = FM:
 GSD     mean    SE   df lower CL upper CL
 ZZ/ZW   14.5 4.776 1.67   -10.43     39.4
 XX/XY   13.8 4.096 0.90   -54.38     82.0

TRN = MF:
 GSD     mean    SE   df lower CL upper CL
 ZZ/ZW   22.5 4.830 2.26     3.83     41.1
 XX/XY   19.2 0.112 1.26    18.33     20.1

Pairwise comparisons:

TRN = FM:
 contrast          estimate   SE    df t-value p-value  lower CL upper CL
 (ZZ/ZW) - (XX/XY)    0.699 2.29 15.60   0.306  0.7640     -4.16     5.56

TRN = MF:
 contrast          estimate   SE    df t-value p-value  lower CL upper CL
 (ZZ/ZW) - (XX/XY)    3.247 4.76  2.29   0.682  0.5578    -14.97    21.46


Reptiles:

λ = 0.90

PGLS coefficients:
                 estimate        SE    t-value p-value
Intercept       20.573510  15.42636  1.3336594  0.1851
 - XX/XY         2.868505  18.78891  0.1526701  0.8789
 - FMF           5.827391  21.08506  0.2763753  0.7828
 - MF            4.510098  16.83001  0.2679795  0.7892
 - XX/XY × FMF  -8.129042  19.02504 -0.4272811  0.6700
 - XX/XY × MF   -6.833035  18.08531 -0.3778224  0.7063

Estimated group means:

TRN = FM:
 GSD     mean   SE   df lower CL upper CL
 ZZ/ZW   20.6 15.4 72.3   -10.18     51.3
 XX/XY   23.4 12.7 72.3    -1.91     48.8

TRN = FMF:
 GSD     mean   SE   df lower CL upper CL
 ZZ/ZW   26.4 15.2 30.5    -4.62     57.4
 XX/XY   21.1 15.2 30.5    -9.95     52.2

TRN = MF:
 GSD     mean   SE   df lower CL upper CL
 ZZ/ZW   25.1 10.6 15.6     2.60     47.6
 XX/XY   21.1  7.2 15.6     5.82     36.4

Pairwise comparisons:

TRN = FM:
 contrast          estimate    SE   df t-value p-value  lower CL upper CL
 (ZZ/ZW) - (XX/XY)    -2.87 18.79 72.3  -0.153  0.8791   -40.320     34.6

TRN = FMF:
 contrast          estimate    SE   df t-value p-value  lower CL upper CL
 (ZZ/ZW) - (XX/XY)     5.26  2.99 30.5   1.760  0.0884    -0.838     11.4

TRN = MF:
 contrast          estimate    SE   df t-value p-value  lower CL upper CL
 (ZZ/ZW) - (XX/XY)     3.96 12.12 15.6   0.327  0.7479   -21.775     29.7


Maximum temperature of warmest month (BIO5) maximum over geographical range

Amphibians:

λ = 0.77

PGLS coefficients:
               estimate        SE   t-value p-value
Intercept      36.29363  4.340430  8.361760  0.0000
 - XX/XY       -2.07507  1.414022 -1.467492  0.1457
 - MF          -2.61005  4.798406 -0.543942  0.5878
 - XX/XY × MF   3.13027  6.253517  0.500562  0.6179

Estimated group means:

TRN = FM:
 GSD     mean      SE   df lower CL upper CL
 ZZ/ZW  36.29 4.34043 6.73    25.95    46.64
 XX/XY  34.22 3.84841 9.28    25.55    42.88

TRN = MF:
 GSD     mean      SE   df lower CL upper CL
 ZZ/ZW  33.68 6.38253 3.02    13.45    53.92
 XX/XY  34.74 0.01191 1.97    34.69    34.79

Pairwise comparisons:

TRN = FM:
 contrast          estimate   SE   df t-value p-value  lower CL upper CL
 (ZZ/ZW) - (XX/XY)     2.08 1.41 6.13   1.467  0.1916     -1.37     5.52

TRN = MF:
 contrast          estimate   SE   df t-value p-value  lower CL upper CL
 (ZZ/ZW) - (XX/XY)    -1.06 6.37 3.01  -0.166  0.8790         -21.29    19.17


Reptiles:

λ = 0.90

PGLS coefficients:
                estimate        SE    t-value p-value
Intercept       36.26378  12.57312  2.8842303  0.0047
 - XX/XY         0.16417  15.30197  0.0107288  0.9915
 - FMF          -0.50024  15.28171 -0.0327348  0.9739
 - MF            3.57446  13.56997  0.2634095  0.7927
 - XX/XY × FMF   1.98657  15.40959  0.1289177  0.8977
 - XX/XY × MF   -7.71686  14.62278 -0.5277291  0.5988

Estimated group means:

TRN = FM:
 GSD     mean    SE    df lower CL upper CL
 ZZ/ZW   36.3 12.57  8.92     7.78     64.7
 XX/XY   36.4 10.35 11.23    13.71     59.1

TRN = FMF:
 GSD     mean    SE    df lower CL upper CL
 ZZ/ZW   35.8  9.36  7.88    14.11     57.4
 XX/XY   37.9  9.39  7.99    16.26     59.6

TRN = MF:
 GSD     mean    SE    df lower CL upper CL
 ZZ/ZW   39.8  8.29  5.74    19.32     60.4
 XX/XY   32.3  5.65  5.14    17.88     46.7

Pairwise comparisons:

TRN = FM:
 contrast          estimate    SE    df t-value p-value  lower CL upper CL
 (ZZ/ZW) - (XX/XY)   -0.164 15.30  9.84  -0.011  0.9917    -34.34    34.01

TRN = FMF:
 contrast          estimate    SE    df t-value p-value  lower CL upper CL
 (ZZ/ZW) - (XX/XY)   -2.151  1.82 29.78  -1.183  0.2462     -5.86     1.56

TRN = MF:
 contrast          estimate    SE    df t-value p-value  lower CL upper CL
 (ZZ/ZW) - (XX/XY)    7.553  9.50  5.60   0.795  0.4588    -16.09    31.19


Maximum temperature of warmest month (BIO5) median over geographical range

Amphibians:

λ = 0.72

PGLS coefficients:
                estimate        SE   t-value p-value
Intercept      30.341296  3.065305  9.898296  0.0000
 - XX/XY       -3.449839  1.248354 -2.763511  0.0069
 - MF          -0.077013  2.874766 -0.026789  0.9787
 - XX/XY × MF   5.656983  3.035095  1.863857  0.0656

Estimated group means:

TRN = FM:
 GSD     mean   SE   df lower CL upper CL
 ZZ/ZW   30.3 3.07 5.24     22.6     38.1
 XX/XY   26.9 3.48 5.46     18.2     35.6

TRN = MF:
 GSD     mean   SE   df lower CL upper CL
 ZZ/ZW   30.3 3.41 2.94     19.3     41.2
 XX/XY   32.5 1.46 1.87     25.7     39.2

Pairwise comparisons:

TRN = FM:
 contrast          estimate   SE    df t-value p-value  lower CL upper CL
 (ZZ/ZW) - (XX/XY)     3.45 1.25 23.80   2.764  0.0109     0.872     6.03

TRN = MF:
 contrast          estimate   SE    df t-value p-value  lower CL upper CL
 (ZZ/ZW) - (XX/XY)    -2.21 2.50  2.22  -0.883  0.4623   -11.996     7.58


Reptiles:

λ = 0.87

PGLS coefficients:
                 estimate        SE   t-value p-value
Intercept       30.468860  8.272948  3.682951  0.0004
 - XX/XY         0.653620 10.151703  0.064385  0.9488
 - FMF           2.621919  9.995632  0.262306  0.7936
 - MF            6.594421  9.815260  0.671854  0.5031
 - XX/XY × FMF  -1.394887 10.237606 -0.136251  0.8919
 - XX/XY × MF   -9.835511 10.418629 -0.944031  0.3473

Estimated group means:

TRN = FM:
 GSD     mean   SE   df lower CL upper CL
 ZZ/ZW   30.5 8.27 5.96     10.2     50.7
 XX/XY   31.1 6.94 8.15     15.2     47.1

TRN = FMF:
 GSD     mean   SE   df lower CL upper CL
 ZZ/ZW   33.1 6.06 5.93     18.2     48.0
 XX/XY   32.3 6.08 6.05     17.5     47.2

TRN = MF:
 GSD     mean   SE   df lower CL upper CL
 ZZ/ZW   37.1 7.20 4.57     18.0     56.1
 XX/XY   27.9 4.84 3.89     14.3     41.5

Pairwise comparisons:

TRN = FM:
 contrast          estimate    SE    df t-value p-value  lower CL upper CL
 (ZZ/ZW) - (XX/XY)   -0.654 10.15  6.82  -0.064  0.9505    -24.79    23.48

TRN = FMF:
 contrast          estimate    SE    df t-value p-value  lower CL upper CL
 (ZZ/ZW) - (XX/XY)    0.741  1.32 29.73   0.560  0.5796     -1.96     3.45

TRN = MF:
 contrast          estimate    SE    df t-value p-value  lower CL upper CL
 (ZZ/ZW) - (XX/XY)    9.182  8.22  4.41   1.117  0.3211    -12.82    31.19


Mean temperature of wettest quarter (BIO8) mean over geographical range

Amphibians:

λ = 0.95

PGLS coefficients:
                estimate        SE    t-value p-value
Intercept      19.290423  7.045443  2.7379999  0.0074
 - XX/XY       -2.685670  1.351735 -1.9868315  0.0499
 - MF           0.665808  4.324285  0.1539694  0.8780
 - XX/XY × MF   8.603930  6.569345  1.3097089  0.1936

Estimated group means:

TRN = FM:
 GSD     mean   SE   df lower CL upper CL
 ZZ/ZW   19.3 7.05 8.49    3.206     35.4
 XX/XY   16.6 7.34 7.57   -0.483     33.7

TRN = MF:
 GSD     mean   SE   df lower CL upper CL
 ZZ/ZW   20.0 8.69 2.34  -12.663     52.6
 XX/XY   25.9 2.82 1.82   12.536     39.2

Pairwise comparisons:

TRN = FM:
 contrast          estimate   SE    df t-value p-value  lower CL upper CL
 (ZZ/ZW) - (XX/XY)     2.69 1.35 21.47   1.987  0.0598    -0.122     5.49

TRN = MF:
 contrast          estimate   SE    df t-value p-value  lower CL upper CL
 (ZZ/ZW) - (XX/XY)    -5.92 6.21  1.39  -0.953  0.4762   -47.559    35.72


Reptiles:

λ = 0.91

PGLS coefficients:

                  estimate        SE    t-value p-value
Intercept        15.172536  12.07859  1.2561511  0.2118
 - XX/XY          3.041493  14.68758  0.2070793  0.8363
 - FMF           11.383741  15.55228  0.7319662  0.4658
 - MF            11.644289  19.39430  0.6003975  0.5495
 - XX/XY × FMF   -6.855126  14.82295 -0.4624670  0.6447
 - XX/XY × MF   -12.937621  20.12489 -0.6428666  0.5217

Estimated group means:

TRN = FM:
 GSD     mean    SE    df lower CL upper CL
 ZZ/ZW   15.2 12.08  8.31  -12.503     42.8
 XX/XY   18.2  9.92 10.51   -3.750     40.2

TRN = FMF:
 GSD     mean    SE    df lower CL upper CL
 ZZ/ZW   26.6 10.45  5.90    0.895     52.2
 XX/XY   22.7 10.47  5.97   -2.902     48.4

TRN = MF:
 GSD     mean    SE    df lower CL upper CL
 ZZ/ZW   26.8 17.85  5.39  -18.093     71.7
 XX/XY   16.9 12.18  4.81  -14.759     48.6

Pairwise comparisons:

TRN = FM:
 contrast          estimate   SE    df t-value p-value  lower CL upper CL
 (ZZ/ZW) - (XX/XY)    -3.04 14.7  9.18  -0.207  0.8405   -36.169    30.09

TRN = FMF:
 contrast          estimate   SE    df t-value p-value  lower CL upper CL
 (ZZ/ZW) - (XX/XY)     3.81  2.0 27.41   1.908  0.0669    -0.284     7.91

TRN = MF:
 contrast          estimate   SE    df t-value p-value  lower CL upper CL
 (ZZ/ZW) - (XX/XY)     9.90 20.5  5.26   0.484  0.6479   -41.913    61.70


Mean temperature of wettest quarter (BIO8) minimum over geographical range

Amphibians:

λ = 0.87

PGLS coefficients:
                estimate        SE    t-value p-value
Intercept       1.975796 11.793469  0.1675331  0.8673
 - XX/XY       -2.783924  3.429271 -0.8118122  0.4190
 - MF           8.024638  8.906986  0.9009375  0.3700
 - XX/XY × MF   7.469485  3.515264  2.1248719  0.0363

Estimated group means:

TRN = FM:
 GSD     mean    SE   df lower CL upper CL
 ZZ/ZW  1.976 11.79 8.09   -25.17     29.1
 XX/XY -0.808  9.72 8.63   -22.93     21.3

TRN = MF:
 GSD     mean    SE   df lower CL upper CL
 ZZ/ZW 10.000  4.40 2.95    -4.15     24.2
 XX/XY 14.686  1.58 1.91     7.59     21.8

Pairwise comparisons:

TRN = FM:
 contrast          estimate   SE   df t-value p-value  lower CL upper CL
 (ZZ/ZW) - (XX/XY)     2.78 3.43 3.80   0.812  0.4647     -6.94     12.5

TRN = MF:
 contrast          estimate   SE   df t-value p-value  lower CL upper CL
 (ZZ/ZW) - (XX/XY)    -4.69 3.16 1.76  -1.484  0.2916    -20.17     10.8


Reptiles:

λ = 0.90

PGLS coefficients:
                 estimate        SE    t-value p-value
Intercept        3.894658  17.11914  0.2275032  0.8205
 - XX/XY        -3.216155  20.85066 -0.1542471  0.8777
 - FMF          14.976999  26.62442  0.5625285  0.5749
 - MF            4.638560  25.74456  0.1801763  0.8574
 - XX/XY × FMF  -3.587468  21.26710 -0.1686863  0.8664
 - XX/XY × MF   -0.482199  26.70640 -0.0180556  0.9856

Estimated group means:

TRN = FM:
 GSD     mean   SE   df lower CL upper CL
 ZZ/ZW  3.895 17.1 72.0    -30.2     38.0
 XX/XY  0.679 14.1 72.0    -27.5     28.8

TRN = FMF:
 GSD     mean   SE   df lower CL upper CL
 ZZ/ZW 18.872 21.3 30.1    -24.6     62.4
 XX/XY 12.068 21.4 30.1    -31.5     55.7

TRN = MF:
 GSD     mean   SE   df lower CL upper CL
 ZZ/ZW  8.533 23.0 15.6    -40.4     57.5
 XX/XY  4.835 15.7 15.6    -28.5     38.1

Pairwise comparisons:

TRN = FM:
 contrast          estimate    SE   df t-value p-value  lower CL upper CL
 (ZZ/ZW) - (XX/XY)     3.22 20.85 72.0   0.154  0.8778    -38.35     44.8

TRN = FMF:
 contrast          estimate    SE   df t-value p-value  lower CL upper CL
 (ZZ/ZW) - (XX/XY)     6.80  4.19 30.1   1.625  0.1147     -1.75     15.4

TRN = MF:
 contrast          estimate    SE   df t-value p-value  lower CL upper CL
 (ZZ/ZW) - (XX/XY)     3.70 26.38 15.6   0.140  0.8903    -52.34     59.7


Mean temperature of wettest quarter (BIO8) maximum over geographical range

Amphibians:

λ = 0.75

PGLS coefficients:
                estimate        SE   t-value p-value
Intercept      27.215017  3.608735  7.541429  0.0000
 - XX/XY       -3.468411  1.211768 -2.862274  0.0052
 - MF          -5.946518  5.670249 -1.048723  0.2971
 - XX/XY × MF  10.383292  6.500751  1.597245  0.1137

Estimated group means:

TRN = FM:
 GSD     mean   SE   df lower CL upper CL
 ZZ/ZW   27.2 3.61 6.04    18.40     36.0
 XX/XY   23.7 3.53 6.68    15.31     32.2

TRN = MF:
 GSD     mean   SE   df lower CL upper CL
 ZZ/ZW   21.3 7.25 3.17    -1.13     43.7
 XX/XY   28.2 1.18 1.90    22.84     33.5

Pairwise comparisons:

TRN = FM:
 contrast          estimate   SE   df t-value p-value  lower CL upper CL
 (ZZ/ZW) - (XX/XY)     3.47 1.21 41.6   2.862  0.0066      1.02     5.91

TRN = MF:
 contrast          estimate   SE   df t-value p-value  lower CL upper CL
 (ZZ/ZW) - (XX/XY)    -6.91 6.38  2.7  -1.083  0.3659    -28.59    14.76


Reptiles:

λ = 0.88

PGLS coefficients:
                  estimate        SE    t-value p-value
Intercept        21.914714  12.55416  1.7456133  0.0837
 - XX/XY          6.063693  15.37062  0.3944989  0.6940
 - FMF            6.587289  13.24202  0.4974534  0.6199
 - MF             9.818693  14.65537  0.6699724  0.5043
 - XX/XY × FMF   -5.179847  15.40599 -0.3362230  0.7374
 - XX/XY × MF   -13.733202  15.56746 -0.8821739  0.3797


Estimated group means:

TRN = FM:
 GSD     mean    SE    df lower CL upper CL
 ZZ/ZW   21.9 12.55  9.17    -6.40     50.2
 XX/XY   28.0 10.47 12.15     5.19     50.8

TRN = FMF:
 GSD     mean    SE    df lower CL upper CL
 ZZ/ZW   28.5  4.91  7.94    17.15     39.9
 XX/XY   29.4  4.93  8.07    18.03     40.7

TRN = MF:
 GSD     mean    SE    df lower CL upper CL
 ZZ/ZW   31.7 10.51  6.13     6.14     57.3
 XX/XY   24.1  7.09  5.37     6.21     41.9

Pairwise comparisons:

TRN = FM:
 contrast          estimate    SE    df t-value p-value  lower CL upper CL
 (ZZ/ZW) - (XX/XY)   -6.064 15.37 10.35  -0.394  0.7012    -40.16    28.03

TRN = FMF:
 contrast          estimate    SE    df t-value p-value  lower CL upper CL
 (ZZ/ZW) - (XX/XY)   -0.884  1.04 29.48  -0.847  0.4037     -3.02     1.25

TRN = MF:
 contrast          estimate    SE    df t-value p-value  lower CL upper CL
 (ZZ/ZW) - (XX/XY)    7.670 12.01  5.96   0.639  0.5469    -21.77    37.11


Mean temperature of wettest quarter (BIO8) median over geographical range

Amphibians:

λ = 0.88

PGLS coefficients:
                estimate        SE   t-value p-value
Intercept      19.921231  6.155754  3.236197  0.0017
 - XX/XY       -3.085889  1.472376 -2.095856  0.0389
 - MF          -1.324296  4.245585 -0.311923  0.7558
 - XX/XY × MF  10.297439  4.708013  2.187216  0.0313

Estimated group means:

TRN = FM:
 GSD     mean   SE   df lower CL upper CL
 ZZ/ZW   19.9 6.16 5.47    4.500     35.3
 XX/XY   16.8 6.18 4.38    0.247     33.4

TRN = MF:
 GSD     mean   SE   df lower CL upper CL
 ZZ/ZW   18.6 6.74 2.26   -7.466     44.7
 XX/XY   25.8 2.91 1.67   10.585     41.0

Pairwise comparisons:

TRN = FM:
 contrast          estimate   SE    df t-value p-value  lower CL upper CL
 (ZZ/ZW) - (XX/XY)     3.09 1.47 71.99   2.096  0.0396     0.151     6.02

TRN = MF:
 contrast          estimate   SE    df t-value p-value  lower CL upper CL
 (ZZ/ZW) - (XX/XY)    -7.21 4.45  1.42  -1.622  0.2932   -36.150    21.73


Reptiles:

λ = 0.89

PGLS coefficients:
                  estimate        SE    t-value p-value
Intercept        15.460719  12.02163  1.2860756  0.2012
 - XX/XY          4.074522  14.67515  0.2776477  0.7818
 - FMF           11.517870  15.30842  0.7523881  0.4535
 - MF            13.735661  18.68773  0.7350097  0.4639
 - XX/XY × FMF   -8.154066  14.81920 -0.5502365  0.5833
 - XX/XY × MF   -15.998460  19.41519 -0.8240178  0.4118

Estimated group means:

TRN = FM:
 GSD     mean    SE    df lower CL upper CL
 ZZ/ZW   15.5 12.02  7.99   -12.27     43.2
 XX/XY   19.5  9.96 10.45    -2.53     41.6

TRN = FMF:
 GSD     mean    SE    df lower CL upper CL
 ZZ/ZW   27.0 10.12  6.01     2.22     51.7
 XX/XY   22.9 10.15  6.10    -1.85     47.6

TRN = MF:
 GSD     mean    SE    df lower CL upper CL
 ZZ/ZW   29.2 16.96  5.55   -13.12     71.5
 XX/XY   17.3 11.49  4.87   -12.51     47.1

Pairwise comparisons:

TRN = FM:
 contrast          estimate    SE    df t-value p-value  lower CL upper CL
 (ZZ/ZW) - (XX/XY)    -4.07 14.68  8.96  -0.278  0.7876   -37.293     29.1

TRN = FMF:
 contrast          estimate    SE    df t-value p-value  lower CL upper CL
 (ZZ/ZW) - (XX/XY)     4.08  2.06 27.64   1.979  0.0578    -0.145      8.3

TRN = MF:
 contrast          estimate    SE    df t-value p-value  lower CL upper CL
 (ZZ/ZW) - (XX/XY)    11.92 19.40  5.40   0.615  0.5638   -36.864     60.7


Mean temperature of warmest quarter (BIO10) mean over geographical range

Amphibians:

λ = 0.73

PGLS coefficients:
                estimate        SE   t-value p-value
Intercept      23.588818  2.634454  8.953969  0.0000
 - XX/XY       -3.408761  1.244225 -2.739666  0.0074
 - MF          -0.032803  2.188908 -0.014986  0.9881
 - XX/XY × MF   7.043510  2.341551  3.008053  0.0034

Estimated group means:

TRN = FM:
 GSD     mean   SE   df lower CL upper CL
 ZZ/ZW   23.6 2.63 4.04     16.3     30.9
 XX/XY   20.2 3.32 3.69     10.6     29.7

TRN = MF:
 GSD     mean   SE   df lower CL upper CL
 ZZ/ZW   23.6 2.39 2.69     15.4     31.7
 XX/XY   27.2 1.31 1.74     20.6     33.7

Pairwise comparisons:

TRN = FM:
 contrast          estimate   SE   df t-value p-value  lower CL upper CL
 (ZZ/ZW) - (XX/XY)     3.41 1.24 7.31   2.740  0.0277     0.492     6.33

TRN = MF:
 contrast          estimate   SE   df t-value p-value  lower CL upper CL
 (ZZ/ZW) - (XX/XY)    -3.63 1.63 2.30  -2.234  0.1381    -9.820     2.55


Reptiles:

λ = 0.91

PGLS coefficients:
                 estimate        SE    t-value p-value
Intercept       22.986131  8.639736  2.6605129  0.0090
 - XX/XY         0.685202 10.497204  0.0652747  0.9481
 - FMF           4.098135  9.754342  0.4201345  0.6752
 - MF            5.626121 12.365428  0.4549880  0.6500
 - XX/XY × FMF  -2.120673 10.539454 -0.2012128  0.8409
 - XX/XY × MF   -7.918631 12.809517 -0.6181834  0.5378

Estimated group means:

TRN = FM:
 GSD     mean    SE   df lower CL upper CL
 ZZ/ZW   23.0  8.64 7.30    2.724     43.2
 XX/XY   23.7  7.08 9.12    7.677     39.7

TRN = FMF:
 GSD     mean    SE   df lower CL upper CL
 ZZ/ZW   27.1  5.00 6.64   15.132     39.0
 XX/XY   25.6  5.01 6.72   13.702     37.6

TRN = MF:
 GSD     mean    SE   df lower CL upper CL
 ZZ/ZW   28.6 10.80 4.91    0.695     56.5
 XX/XY   21.4  7.38 4.39    1.597     41.2

Pairwise comparisons:

TRN = FM:
 contrast          estimate     SE    df t-value p-value  lower CL upper CL
 (ZZ/ZW) - (XX/XY)   -0.685 10.497  8.02  -0.065  0.9496   -24.882    23.51

TRN = FMF:
 contrast          estimate     SE    df t-value p-value  lower CL upper CL
 (ZZ/ZW) - (XX/XY)    1.435  0.943 29.76   1.523  0.1384    -0.491     3.36

TRN = MF:
 contrast          estimate     SE    df t-value p-value  lower CL upper CL
 (ZZ/ZW) - (XX/XY)    7.233 12.374  4.79   0.585  0.5853   -25.004    39.47


Mean temperature of warmest quarter (BIO10) minimum over geographical range

Amphibians:

λ = 0.74

PGLS coefficients:
                estimate        SE    t-value p-value
Intercept       8.754197  6.285405  1.3927816  0.1671
 - XX/XY       -1.117708  2.149555 -0.5199720  0.6043
 - MF           8.841312  5.638933  1.5679052  0.1204
 - XX/XY × MF  -1.503950  6.040336 -0.2489844  0.8039

Estimated group means:

TRN = FM:
 GSD     mean   SE   df lower CL upper CL
 ZZ/ZW   8.75 6.29 3.28   -10.31     27.8
 XX/XY   7.64 5.67 2.19   -14.89     30.2

TRN = MF:
 GSD     mean   SE   df lower CL upper CL
 ZZ/ZW  17.60 6.79 2.42    -7.25     42.4
 XX/XY  14.97 1.17 1.52     8.10     21.8

Pairwise comparisons:

TRN = FM:
 contrast          estimate   SE    df t-value p-value  lower CL upper CL
 (ZZ/ZW) - (XX/XY)     1.12 2.15 12.00   0.520  0.6125     -3.57      5.8

TRN = MF:
 contrast          estimate   SE    df t-value p-value  lower CL upper CL
 (ZZ/ZW) - (XX/XY)     2.62 5.95  2.26   0.441  0.6979    -20.33     25.6


Reptiles:

λ = 0.90

PGLS coefficients:
                 estimate        SE    t-value p-value
Intercept       13.862830  14.28081  0.9707311  0.3339
 - XX/XY         2.327036  17.39366  0.1337864  0.8938
 - FMF           7.197189  20.28593  0.3547872  0.7234
 - MF            4.933004  16.44897  0.2998975  0.7648
 - XX/XY × FMF  -8.484617  17.64751 -0.4807827  0.6317
 - XX/XY × MF   -5.982123  17.41901 -0.3434249  0.7320

Estimated group means:

TRN = FM:
 GSD     mean    SE   df lower CL upper CL
 ZZ/ZW   13.9 14.28 70.9   -14.61     42.3
 XX/XY   16.2 11.77 70.9    -7.29     39.7

TRN = FMF:
 GSD     mean    SE   df lower CL upper CL
 ZZ/ZW   21.1 15.17 28.7    -9.98     52.1
 XX/XY   14.9 15.21 28.7   -16.21     46.0

TRN = MF:
 GSD     mean    SE   df lower CL upper CL
 ZZ/ZW   18.8 11.59 15.6    -5.82     43.4
 XX/XY   15.1  7.88 15.6    -1.61     31.9

Pairwise comparisons:

TRN = FM:
 contrast          estimate    SE   df t-value p-value  lower CL upper CL
 (ZZ/ZW) - (XX/XY)    -2.33 17.39 70.9  -0.134  0.8940   -37.010     32.4

TRN = FMF:
 contrast          estimate    SE   df t-value p-value  lower CL upper CL
 (ZZ/ZW) - (XX/XY)     6.16  2.98 28.7   2.065  0.0481     0.055     12.3

TRN = MF:
 contrast          estimate    SE   df t-value p-value  lower CL upper CL
 (ZZ/ZW) - (XX/XY)     3.66 13.27 15.6   0.276  0.7865   -24.527     31.8


Mean temperature of warmest quarter (BIO10) maximum over geographical range

Amphibians:

λ = 0.69

PGLS coefficients:
                estimate        SE   t-value p-value
Intercept      28.896167  2.807569 10.292238  0.0000
 - XX/XY       -1.966216  1.035482 -1.898841  0.0608
 - MF          -2.088373  3.311804 -0.630585  0.5299
 - XX/XY × MF   4.562623  3.891842  1.172356  0.2441

Estimated group means:

TRN = FM:
 GSD     mean   SE   df lower CL upper CL
 ZZ/ZW   28.9 2.81 7.13     22.3     35.5
 XX/XY   26.9 2.63 8.56     20.9     32.9

TRN = MF:
 GSD     mean   SE   df lower CL upper CL
 ZZ/ZW   26.8 4.07 3.22     14.3     39.3
 XX/XY   29.4 0.37 1.98     27.8     31.0

Pairwise comparisons:

TRN = FM:
 contrast          estimate   SE    df t-value p-value  lower CL upper CL
 (ZZ/ZW) - (XX/XY)     1.97 1.04 21.50   1.899  0.0711    -0.184     4.12

TRN = MF:
 contrast          estimate   SE    df t-value p-value  lower CL upper CL
 (ZZ/ZW) - (XX/XY)    -2.60 3.81  2.94  -0.682  0.5451   -14.852     9.66


Reptiles:

λ = 0.93

PGLS coefficients:
                 estimate        SE    t-value p-value
Intercept       28.284368  11.85167  2.3865296  0.0188
 - XX/XY         0.160439  14.33895  0.0111891  0.9911
 - FMF           0.668899  13.06926  0.0511811  0.9593
 - MF            3.411028  13.92438  0.2449680  0.8070
 - XX/XY × FMF   1.648413  14.37864  0.1146432  0.9089
 - XX/XY × MF   -6.617002  14.57975 -0.4538488  0.6509

Estimated group means:

TRN = FM:
 GSD     mean    SE    df lower CL upper CL
 ZZ/ZW   28.3 11.85 11.59     2.36     54.2
 XX/XY   28.4  9.62 13.73     7.77     49.1

TRN = FMF:
 GSD     mean    SE    df lower CL upper CL
 ZZ/ZW   29.0  6.16 10.46    15.31     42.6
 XX/XY   30.8  6.17 10.54    17.11     44.4

TRN = MF:
 GSD     mean    SE    df lower CL upper CL
 ZZ/ZW   31.7 10.17  6.54     7.29     56.1
 XX/XY   25.2  7.00  6.06     8.15     42.3

Pairwise comparisons:

TRN = FM:
 contrast          estimate    SE    df t-value p-value  lower CL upper CL
 (ZZ/ZW) - (XX/XY)    -0.16 14.34 12.45  -0.011  0.9912    -31.28   30.957

TRN = FMF:
 contrast          estimate    SE    df t-value p-value  lower CL upper CL
 (ZZ/ZW) - (XX/XY)    -1.81  1.07 30.08  -1.694  0.1006     -3.99    0.371

TRN = MF:
 contrast          estimate    SE    df t-value p-value  lower CL upper CL
 (ZZ/ZW) - (XX/XY)     6.46 11.67  6.43   0.553  0.5989    -21.66   34.570


Mean temperature of warmest quarter (BIO10) median over geographical range

Amphibians:

λ = 0.68

PGLS coefficients:
                estimate        SE   t-value p-value
Intercept      23.946994  2.545737  9.406705  0.0000
 - XX/XY       -3.567086  1.262187 -2.826116  0.0058
 - MF          -0.450100  2.251601 -0.199902  0.8420
 - XX/XY × MF   7.770675  2.249088  3.455034  0.0008

Estimated group means:

TRN = FM:
 GSD     mean   SE   df lower CL upper CL
 ZZ/ZW   23.9 2.55 3.72     16.7     31.2
 XX/XY   20.4 3.17 3.17     10.6     30.2

TRN = MF:
 GSD     mean   SE   df lower CL upper CL
 ZZ/ZW   23.5 2.34 2.61     15.4     31.6
 XX/XY   27.7 1.66 1.72     19.3     36.1

Pairwise comparisons:

TRN = FM:
 contrast          estimate   SE   df t-value p-value  lower CL upper CL
 (ZZ/ZW) - (XX/XY)     3.57 1.26 9.12   2.826  0.0196     0.718    6.417

TRN = MF:
 contrast          estimate   SE   df t-value p-value  lower CL upper CL
 (ZZ/ZW) - (XX/XY)    -4.20 1.61 3.89  -2.617  0.0606    -8.713    0.305


Reptiles:

λ = 0.91

PGLS coefficients:
                 estimate        SE    t-value p-value
Intercept       23.215919  9.079329  2.5570081  0.0120
 - XX/XY         0.666906 11.022647  0.0605032  0.9519
 - FMF           4.197376 10.185431  0.4120960  0.6811
 - MF            6.065911 12.732404  0.4764152  0.6348
 - XX/XY × FMF  -2.259129 11.063538 -0.2041959  0.8386
 - XX/XY × MF   -8.408467 13.186474 -0.6376585  0.5251

Estimated group means:

TRN = FM:
 GSD     mean    SE   df lower CL upper CL
 ZZ/ZW   23.2  9.08 7.69     2.13     44.3
 XX/XY   23.9  7.43 9.51     7.21     40.6

TRN = FMF:
 GSD     mean    SE   df lower CL upper CL
 ZZ/ZW   27.4  5.11 7.26    15.41     39.4
 XX/XY   25.8  5.12 7.34    13.82     37.8

TRN = MF:
 GSD     mean    SE   df lower CL upper CL
 ZZ/ZW   29.3 10.99 5.15     1.28     57.3
 XX/XY   21.5  7.52 4.64     1.75     41.3

Pairwise comparisons:

TRN = FM:
 contrast          estimate    SE    df t-value p-value  lower CL upper CL
 (ZZ/ZW) - (XX/XY)   -0.667 11.02  8.42  -0.061  0.9532   -25.868    24.53

TRN = FMF:
 contrast          estimate    SE    df t-value p-value  lower CL upper CL
 (ZZ/ZW) - (XX/XY)    1.592  0.95 29.98   1.675  0.1042    -0.349     3.53

TRN = MF:
 contrast          estimate    SE    df t-value p-value  lower CL upper CL
 (ZZ/ZW) - (XX/XY)    7.742 12.60  5.03   0.615  0.5655   -24.576    40.06
```
