## Supplementary Table 2 for "Interplay of genotypic and thermal sex determination shapes climatic distribution in herpetofauna"

**Supplementary Table 2. Pairwise Pearson correlation coefficients between each of the 16 WorldClim variables (columns) and each of the 20 SexClim variables (rows; see footnotes for the explanations of row numbers) for A) amphibians and B) reptiles.** Correlations in which the two climatic variables refer to the same aspect of climatic distribution (i.e. mean, median, minimum, or maximum temperature across the geographical distribution range) are highlighted with black bold font and thick borders; μ is the mean of the latter correlations for each WorldClim variable.

| **A)** | BIO1 mean | BIO5 mean | BIO8 mean | BIO10 mean | BIO1 median | BIO5 median | BIO8 median | BIO10 median | BIO1 min | BIO5 min | BIO8 min | BIO10 min | BIO1 max | BIO5 max | BIO8 max | BIO10 max |
| --- | --- | --- | --- | --- | --- | --- | --- | --- | --- | --- | --- | --- | --- | --- | --- | --- |
| 1 | **0.794** | **0.833** | **0.688** | **0.877** | 0.785 | 0.820 | 0.701 | 0.863 | 0.558 | 0.462 | 0.524 | 0.526 | 0.801 | 0.608 | 0.594 | 0.682 |
| 2 | **0.748** | **0.880** | **0.629** | **0.905** | 0.738 | 0.872 | 0.646 | 0.895 | 0.470 | 0.421 | 0.446 | 0.475 | 0.803 | 0.673 | 0.562 | 0.747 |
| 3 | **0.738** | **0.673** | **0.656** | **0.732** | 0.732 | 0.657 | 0.662 | 0.717 | 0.566 | 0.431 | 0.530 | 0.496 | 0.693 | 0.466 | 0.542 | 0.531 |
| 4 | **0.803** | **0.866** | **0.705** | **0.907** | 0.795 | 0.857 | 0.717 | 0.896 | 0.543 | 0.463 | 0.518 | 0.522 | 0.816 | 0.643 | 0.618 | 0.721 |
| 5 | **0.742** | **0.925** | **0.678** | **0.947** | 0.734 | 0.923 | 0.696 | 0.943 | 0.444 | 0.424 | 0.435 | 0.471 | 0.817 | 0.714 | 0.634 | 0.796 |
| 6 | **0.751** | **0.669** | **0.670** | **0.734** | 0.746 | 0.654 | 0.673 | 0.718 | 0.553 | 0.406 | 0.537 | 0.472 | 0.702 | 0.474 | 0.552 | 0.544 |
| 7 | 0.806 | 0.845 | 0.698 | 0.894 | **0.798** | **0.835** | **0.711** | **0.884** | 0.560 | 0.467 | 0.526 | 0.531 | 0.810 | 0.615 | 0.603 | 0.690 |
| 8 | 0.757 | 0.884 | 0.643 | 0.914 | **0.749** | **0.878** | **0.660** | **0.907** | 0.475 | 0.424 | 0.456 | 0.478 | 0.806 | 0.672 | 0.572 | 0.747 |
| 9 | 0.750 | 0.670 | 0.666 | 0.737 | **0.747** | **0.656** | **0.671** | **0.725** | 0.564 | 0.427 | 0.537 | 0.491 | 0.694 | 0.460 | 0.542 | 0.526 |
| 10 | 0.792 | 0.869 | 0.671 | 0.905 | **0.786** | **0.863** | **0.684** | **0.899** | 0.529 | 0.465 | 0.499 | 0.518 | 0.796 | 0.644 | 0.578 | 0.715 |
| 11 | 0.747 | 0.922 | 0.690 | 0.950 | **0.741** | **0.923** | **0.708** | **0.950** | 0.444 | 0.422 | 0.444 | 0.469 | 0.814 | 0.708 | 0.641 | 0.791 |
| 12 | 0.762 | 0.664 | 0.679 | 0.736 | **0.759** | **0.651** | **0.681** | **0.725** | 0.551 | 0.401 | 0.544 | 0.465 | 0.701 | 0.467 | 0.551 | 0.537 |
| 13 | 0.359 | 0.200 | 0.242 | 0.235 | 0.344 | 0.196 | 0.232 | 0.222 | **0.763** | **0.793** | **0.595** | **0.809** | 0.103 | -0.223 | 0.005 | -0.149 |
| 14 | 0.482 | 0.345 | 0.313 | 0.391 | 0.467 | 0.342 | 0.304 | 0.378 | **0.831** | **0.879** | **0.662** | **0.904** | 0.227 | -0.121 | 0.056 | -0.034 |
| 15 | 0.359 | 0.200 | 0.242 | 0.235 | 0.344 | 0.196 | 0.232 | 0.222 | **0.763** | **0.793** | **0.595** | **0.809** | 0.103 | -0.223 | 0.005 | -0.149 |
| 16 | 0.482 | 0.345 | 0.313 | 0.391 | 0.467 | 0.342 | 0.304 | 0.378 | **0.831** | **0.879** | **0.662** | **0.904** | 0.227 | -0.121 | 0.056 | -0.034 |
| 17 | 0.548 | 0.847 | 0.480 | 0.807 | 0.538 | 0.834 | 0.514 | 0.791 | 0.109 | 0.078 | 0.092 | 0.114 | **0.797** | **0.892** | **0.610** | **0.917** |
| 18 | 0.589 | 0.833 | 0.547 | 0.810 | 0.578 | 0.815 | 0.580 | 0.790 | 0.178 | 0.114 | 0.154 | 0.165 | **0.826** | **0.865** | **0.661** | **0.896** |
| 19 | 0.548 | 0.847 | 0.480 | 0.807 | 0.538 | 0.834 | 0.514 | 0.791 | 0.109 | 0.078 | 0.092 | 0.114 | **0.797** | **0.892** | **0.610** | **0.917** |
| 20 | 0.589 | 0.833 | 0.547 | 0.810 | 0.578 | 0.815 | 0.580 | 0.790 | 0.178 | 0.114 | 0.154 | 0.165 | **0.826** | **0.865** | **0.661** | **0.896** |
| **μ** | **0.729** | **0.655** | **0.619** | **0.718** | **0.724** | **0.653** | **0.617** | **0.714** | **0.758** | **0.665** | **0.585** | **0.692** | **0.773** | **0.610** | **0.714** | **0.691** |

| **B)** | BIO1 mean | BIO5 mean | BIO8 mean | BIO10 mean | BIO1 median | BIO5 median | BIO8 median | BIO10 median | BIO1 min | BIO5 min | BIO8 min | BIO10 min | BIO1 max | BIO5 max | BIO8 max | BIO10 max |
| --- | --- | --- | --- | --- | --- | --- | --- | --- | --- | --- | --- | --- | --- | --- | --- | --- |
| 1 | **0.794** | **0.833** | **0.688** | **0.877** | 0.785 | 0.820 | 0.701 | 0.863 | 0.558 | 0.462 | 0.524 | 0.526 | 0.801 | 0.608 | 0.594 | 0.682 |
| 2 | **0.748** | **0.880** | **0.629** | **0.905** | 0.738 | 0.872 | 0.646 | 0.895 | 0.470 | 0.421 | 0.446 | 0.475 | 0.803 | 0.673 | 0.562 | 0.747 |
| 3 | **0.738** | **0.673** | **0.656** | **0.732** | 0.732 | 0.657 | 0.662 | 0.717 | 0.566 | 0.431 | 0.530 | 0.496 | 0.693 | 0.466 | 0.542 | 0.531 |
| 4 | **0.803** | **0.866** | **0.705** | **0.907** | 0.795 | 0.857 | 0.717 | 0.896 | 0.543 | 0.463 | 0.518 | 0.522 | 0.816 | 0.643 | 0.618 | 0.721 |
| 5 | **0.742** | **0.925** | **0.678** | **0.947** | 0.734 | 0.923 | 0.696 | 0.943 | 0.444 | 0.424 | 0.435 | 0.471 | 0.817 | 0.714 | 0.634 | 0.796 |
| 6 | **0.751** | **0.669** | **0.670** | **0.734** | 0.746 | 0.654 | 0.673 | 0.718 | 0.553 | 0.406 | 0.537 | 0.472 | 0.702 | 0.474 | 0.552 | 0.544 |
| 7 | 0.806 | 0.845 | 0.698 | 0.894 | **0.798** | **0.835** | **0.711** | **0.884** | 0.560 | 0.467 | 0.526 | 0.531 | 0.810 | 0.615 | 0.603 | 0.690 |
| 8 | 0.757 | 0.884 | 0.643 | 0.914 | **0.749** | **0.878** | **0.660** | **0.907** | 0.475 | 0.424 | 0.456 | 0.478 | 0.806 | 0.672 | 0.572 | 0.747 |
| 9 | 0.750 | 0.670 | 0.666 | 0.737 | **0.747** | **0.656** | **0.671** | **0.725** | 0.564 | 0.427 | 0.537 | 0.491 | 0.694 | 0.460 | 0.542 | 0.526 |
| 10 | 0.792 | 0.869 | 0.671 | 0.905 | **0.786** | **0.863** | **0.684** | **0.899** | 0.529 | 0.465 | 0.499 | 0.518 | 0.796 | 0.644 | 0.578 | 0.715 |
| 11 | 0.747 | 0.922 | 0.690 | 0.950 | **0.741** | **0.923** | **0.708** | **0.950** | 0.444 | 0.422 | 0.444 | 0.469 | 0.814 | 0.708 | 0.641 | 0.791 |
| 12 | 0.762 | 0.664 | 0.679 | 0.736 | **0.759** | **0.651** | **0.681** | **0.725** | 0.551 | 0.401 | 0.544 | 0.465 | 0.701 | 0.467 | 0.551 | 0.537 |
| 13 | 0.359 | 0.200 | 0.242 | 0.235 | 0.344 | 0.196 | 0.232 | 0.222 | **0.763** | **0.793** | **0.595** | **0.809** | 0.103 | -0.223 | 0.005 | -0.149 |
| 14 | 0.482 | 0.345 | 0.313 | 0.391 | 0.467 | 0.342 | 0.304 | 0.378 | **0.831** | **0.879** | **0.662** | **0.904** | 0.227 | -0.121 | 0.056 | -0.034 |
| 15 | 0.359 | 0.200 | 0.242 | 0.235 | 0.344 | 0.196 | 0.232 | 0.222 | **0.763** | **0.793** | **0.595** | **0.809** | 0.103 | -0.223 | 0.005 | -0.149 |
| 16 | 0.482 | 0.345 | 0.313 | 0.391 | 0.467 | 0.342 | 0.304 | 0.378 | **0.831** | **0.879** | **0.662** | **0.904** | 0.227 | -0.121 | 0.056 | -0.034 |
| 17 | 0.548 | 0.847 | 0.480 | 0.807 | 0.538 | 0.834 | 0.514 | 0.791 | 0.109 | 0.078 | 0.092 | 0.114 | **0.797** | **0.892** | **0.610** | **0.917** |
| 18 | 0.589 | 0.833 | 0.547 | 0.810 | 0.578 | 0.815 | 0.580 | 0.790 | 0.178 | 0.114 | 0.154 | 0.165 | **0.826** | **0.865** | **0.661** | **0.896** |
| 19 | 0.548 | 0.847 | 0.480 | 0.807 | 0.538 | 0.834 | 0.514 | 0.791 | 0.109 | 0.078 | 0.092 | 0.114 | **0.797** | **0.892** | **0.610** | **0.917** |
| 20 | 0.589 | 0.833 | 0.547 | 0.810 | 0.578 | 0.815 | 0.580 | 0.790 | 0.178 | 0.114 | 0.154 | 0.165 | **0.826** | **0.865** | **0.661** | **0.896** |
| **μ** | **0.763** | **0.808** | **0.671** | **0.850** | **0.763** | **0.801** | **0.686** | **0.848** | **0.797** | **0.836** | **0.629** | **0.857** | **0.812** | **0.878** | **0.636** | **0.907** |

Row numbers represent SexClim variables as follows (see Metadata for more detailed explanation):

1	Mean_temp_breeding

2	Mean_temp_warmest_breeding_month

3	Mean_temp_coldest_breeding_month

4	Mean_temp_extended_breeding

5	Mean_temp_warmest_extended_breeding_month

6	Mean_temp_coldest_extended_breeding_month

7	Median_temp_breeding

8	Median_temp_warmest_breeding_month

9	Median_temp_coldest_breeding_month

10	Median_temp_extended_breeding

11	Median_temp_warmest_extended_breeding_month

12	Median_temp_coldest_extended_breeding_month

13	Min_temp_coldest_breeding_month

14	Mean_temp_at_coldest_area_breeding

15	Min_temp_coldest_extended_breeding_month

16	Mean_temp_at_coldest_extended_area_breeding

17	Max_temp_warmest_breeding_month

18	Mean_temp_at_warmest_area_breeding

19	Max_temp_warmest_extended_breeding_month

20	Mean_temp_at_warmest_extended_area_breeding
