## Supplementary Table 3 for "Interplay of genotypic and thermal sex determination shapes climatic distribution in herpetofauna"

**Supplementary Table 3. Justification for the clade-specific categorization of temperature reaction norm (TRN) patterns used in the present study, based on the information available in the HerpSexDet database (version 1.1).** Note that we used only those taxonomic families where genotypic sex determination (GSD) occurs in at least one species according to HerpSexDet. To each of these families, we assigned the TRN pattern that was reported for at least two species if no other TRN pattern was reported in the family. If more than one TRN pattern occurred in a family, we applied this rule to smaller phylogenetic clades within the family whenever possible. We assigned species-specific TRN to those GSD species for which their TRN was reported, regardless of the TRN patterns in the rest of the family.

| **Taxonomic families** | **Number of species with TRN data in HerpSexDet** | | | | | **TRN classification used in the present study, with justification** |
| --- | --- | --- | --- | --- | --- | --- |
|  | **FM** | **FMF** | **FMFM** | **MF** | **MFM** |  |
| **Amphibia** |  |  |  |  |  |  |
| ***Anura*** |  |  |  |  |  |  |
| Bufonidae | 1 |  |  |  |  | FM: info on 1 species in the family, accepted because it agrees with all anurans excepting Pipidae. |
| Hylidae | 1 |  |  |  |  | FM: info on 1 species in the family, accepted because it agrees with all anurans excepting Pipidae. |
| Pipidae |  |  |  | 4 |  | MF: info on several hybrids (combinations of 4 species; collated into a single row in HerpSexDet). |
| Ranidae | 5 |  |  |  |  | FM: all species with TRN data in this family have FM pattern. |
| ***Caudata*** |  |  |  |  |  |  |
| Cryptobranchidae | 1 |  |  |  |  | FM: known for *Andrias davidianus* (the only species with known GSD in this family). |
| Hynobiidae |  |  |  | 1 |  | NA: TRN info on 1 species in the family (MF), conflicted by closest relative (Andrias: FM). |
| Salamandridae | 3 |  |  | 1 |  | FM in *Pleurodeles waltl* and MF in *Pleurodeles poireti* (based on TRN reported for these two species). FM for the rest of the family because they belong to the clade with 2 FM species (former Triturus genus). |
| **Reptilia** |  |  |  |  |  |  |
| ***Squamata*** |  |  |  |  |  |  |
| Agamidae | 6 | 6 | 1 | 1 |  | MF in *Pogona vitticeps* (based on TRN reported for this species). NA for 4 other species because FM, MF, FMF, and FMFM types occur in the family, with no apparent phylogenetic grouping. |
| Crotaphytidae |  |  |  |  | 1 | NA: info on 1 species in the family (MFM), conflicted by closest families (Iguanidae: 1 FMF species, Tropiduridae: 1 MFM species). |
| Diplodactylidae |  | 2 |  |  |  | FMF: found in 2 species in this family, we extrapolated this info to the 1 species present in our dataset. |
| Eublepharidae |  | 2 |  |  |  | FMF: found in 2 species in this family, we extrapolated this info for the 2 other species present in our dataset. |
| Gekkonidae |  | 2 |  |  |  | FMF: found in 2 species in this family, we extrapolated this info to the other species present in our dataset. |
| Iguanidae |  | 1 |  |  |  | NA: info on 1 species in the family (FMF), conflicted by closest families (Tropiduridae & Crotaphytidae: 1 MFM species in each family). |
| Lacertidae | 2 |  |  |  |  | FM: found in 2 species in this family, we extrapolated this info to the other species present in our dataset. |
| Phyllodactylidae |  |  |  | 1 |  | NA: info on 1 species in the family (MF?), conflicted by closest families (FMF in Gekkonidae & Eublepharidae). |
| Scincidae | 2 |  |  | 2 |  | 2 FM and 2 MF species occur in 2 separate clades within the family, respectively. We categorized the 10 species with GSD in these 2 clades accordingly. The remaining 10 species that do not fall into these 2 clades are set to NA. |
| Tropiduridae |  |  |  |  | 1 | NA: info on 1 species in the family (MFM), conflicted by closest families (Iguanidae: 1 FMF species, Crotaphytidae: 1 MFM species). |
| Viperidae | 1 |  |  |  |  | NA: info on 1 species in the family, not accepted because no other snake is known to have TRN. |
| ***Testudines*** |  |  |  |  |  |  |
| Emydidae |  |  |  | 27 |  | MF: all species with TRN data in this family have MF pattern. |
| Geoemydidae |  | 3 |  | 12 |  | MF: the closest clade is the Batagur genus, which has 3 MF species. |
| Kinosternidae |  | 9 |  | 4 |  | NA: no direct info; the other 2 genera in the family have FMF (Sternotherus) or both FMF and MF (Kinosternon). The closest family is Dermatemydidae, with MF. |
